# A farnesyl-dependent structural role for CENP-E in expansion of the fibrous corona

**DOI:** 10.1101/2023.04.26.538394

**Authors:** Jingchao Wu, Maximilian W.D. Raas, Paula Sobrevals Alcaraz, Harmjan R. Vos, Eelco C. Tromer, Berend Snel, Geert J.P.L. Kops

## Abstract

Correct chromosome segregation during cell division depends on proper connections between spindle microtubules and kinetochores. During prometaphase, kinetochores are temporarily covered with a dense protein meshwork known as the fibrous corona. Formed by oligomerization of ROD/ZW10/ZWILCH-SPINDLY (RZZ-S) complexes, the fibrous corona promotes spindle assembly, chromosome orientation and spindle checkpoint signaling. The molecular requirements for formation of the fibrous corona are not fully understood. Here we show that the fibrous corona depends on the mitotic kinesin CENP-E, and that poorly expanded fibrous coronas after CENP-E depletion are functionally compromised. This previously unrecognized role for CENP-E does not require its motor activity but instead is driven by farnesyl modification of its C-terminal kinetochore-and microtubule-binding domain. We show that in cells CENP-E interacts with RZZ-S complexes in a farnesyl-dependent manner. CENP-E is recruited to kinetochores following RZZ-S, and - while not required for RZZ-S oligomerization per se - promotes subsequent fibrous corona expansion. Our comparative genomics analyses suggest that the farnesylation motif in CENP-E orthologs emerged alongside the full RZZ-S module in an ancestral lineage close to the fungi-animal split (Obazoa), revealing potential conservation of the mechanisms for fibrous corona formation. Our results show that proper spindle assembly has a potentially conserved non-motor contribution from the kinesin CENP-E through stabilization of the fibrous corona meshwork during its formation.

## Introduction

The cell division cycle culminates in the segregation of duplicated sister chromatids to nascent daughter cells. Segregation is driven by the microtubule-based spindle apparatus (Pavin and Tolic, 2021). Microtubules connect to chromosomes via kinetochores, large protein complexes that assemble on centromeric chromatin and that contain microtubule-binding proteins (Cheeseman, 2014; Musacchio and Desai, 2017). To facilitate spindle assembly, kinetochores transiently expand in early mitosis by forming a voluminous protein meshwork known as the fibrous corona (Kops and Gassmann, 2020). The fibrous corona can make extensive connections to microtubule lattices (Kapoor et al., 2006; Rieder and Alexander, 1990; Sacristan et al., 2018), promotes chromosome orientation on the spindle (Magidson et al., 2015), is a microtubule nucleation platform (Mitchison and Kirschner, 1985; Ris and Witt, 1981; Witt et al., 1980; Wu et al., 2023), and strengthens spindle assembly checkpoint (SAC) signaling (Allan et al., 2020; Rodriguez-Rodriguez et al., 2018; Zhang et al., 2019).

The fibrous corona houses dozens of proteins to perform its various roles in spindle assembly and chromosome segregation (Kops and Gassmann, 2020). It is formed by oligomerization of ROD/ZW10/ZWILCH-SPINDLY (RZZ-S) complexes (Mosalaganti et al., 2017; Pereira et al., 2018; Sacristan et al., 2018), which directly bind the dynein/dynactin microtubule motor complex (Chan et al., 2009; d’Amico et al., 2022; Gama et al., 2017; Griffis et al., 2007; Sacristan et al., 2018; Starr et al., 1998). Dynein/dynactin aid in microtubule capture and subsequent poleward transport of chromosomes along the microtubule lattice to facilitate eventual biorientation (Li et al., 2007; Savoian et al., 2000; Vorozhko et al., 2008; Yang et al., 2007). In addition, the LIC1 subunit of dynein promotes microtubule nucleation from the fibrous corona through interaction with the centrosomal protein pericentrin (Wu et al., 2023). Once kinetochores engage in end-on microtubule connections, dynein/dynactin drive the disassembly of the fibrous corona by ‘stripping’ RZZ-S complexes and associated proteins from kinetochores (Basto et al., 2004; Famulski et al., 2011; Howell et al., 2001; Wojcik et al., 2001).

Other proteins of the fibrous corona include the SAC complex MAD1-MAD2- p31^comet^, the Cyclin B1-CDK1 kinase complex, the CLASP1/2 microtubule dynamics regulators, and the microtubule-binding protein CENP-F (Kops and Gassmann, 2020). The fibrous corona additionally harbors CENP-E (Cooke et al., 1997; Thrower et al., 1996; Yao et al., 1997), a kinesin-7 family microtubule motor that promotes correct orientation of microtubule stubs near the kinetochore, stable microtubule interactions, and congression of polar chromosomes (Kapoor et al., 2006; McEwen et al., 2001; Putkey et al., 2002; Schaar et al., 1997; Sikirzhytski et al., 2018; Wood et al., 1997). CENP-E is a dimer with amino-terminal kinesin domains and carboxy-terminal microtubule-binding domains, separated by a discontinuous ∼230 nm coiled-coil (Kim et al., 2008; Yen et al., 1992). The carboxy terminus of human CENP-E contains a CAAX box (with sequence Cys-Lys-Thr-Gln), which is a signal for farnesyl modification on the motif’s cysteine residue (Ashar et al., 2000).

CENP-E localizes to distinct locations on kinetochores. Besides a pool that localizes to fibrous coronas and is stripped by dynein/dynactin upon end-on microtubule binding (Howell et al., 2001), another pool of CENP-E, presumably bound to the core kinetochore, localizes regardless of whether kinetochores are bound to microtubules, and remains associated with kinetochores in metaphase cells (Weaver et al., 2003; Yen et al., 1991; Yen et al., 1992). While a direct interaction of CENP-E with BUBR1 likely contributes to the latter pool (Chan et al., 1998; Ciossani et al., 2018; Legal et al., 2020; Yao et al., 2000), the mechanism for CENP-E’s localization to the fibrous corona is not clear but appears to involve its farnesylation: Farnesyltransferase inhibitor treatment or mutation of the farnesyl-acceptor cysteine residue impact CENP-E levels most prominently on prometaphase kinetochores and, in cell lines where fibrous coronas can be distinguished, prevent CENP-E from localizing to their crescent shapes (Ciossani et al., 2018; Holland et al., 2015; Schafer-Hales et al., 2007).

Fibrous corona formation in early mitosis is driven by oligomerization of RZZ-S complexes, a process that can be recapitulated by purified components in vitro (Pereira et al., 2018; Raisch et al., 2022; Sacristan et al., 2018). It is initiated by phosphorylation of the beta-propeller domain of ROD by the kinetochore kinase MPS1 and involves conformational transitions in SPINDLY (Raisch et al., 2022; Rodriguez-Rodriguez et al., 2018; Sacristan et al., 2018). These events are likely to be mechanistically connected, as fibrous corona expansion no longer requires MPS1 activity when cells express a SPINDLY mutant mimicking its conformationally ‘active’ state (Sacristan et al., 2018). While certainly required, whether these mechanisms are also sufficient for fibrous corona formation in cells is unknown. Here we show that CENP-E has an important and possibly evolutionary conserved structural role in fibrous corona formation.

## Results

### Fibrous corona formation requires CENP-E

Being a constituent of the fibrous corona (Cooke et al., 1997; Thrower et al., 1996; Yao et al., 1997), we wondered whether CENP-E contributes to fibrous corona formation. The area of the fibrous corona in nocodazole-treated RPE1 cells, marked by the RZZ component ZWILCH (Mosalaganti et al., 2017; Pereira et al., 2018), was substantially reduced when CENP-E was depleted by RNAi (Figure 1A, D). Depletion of CENP-E affected fibrous corona formation to a similar extent as loss of SPINDLY (Figure 1B-D) (Pereira et al., 2018; Sacristan et al., 2018). CENP-E depletion did not affect local levels of its outer-kinetochore receptor BUBR1 (Figure S1A-B) but strongly reduced the fibrous corona-localized SAC complex MAD1-MAD2 (Figure S1C-F).

**Figure 1.**
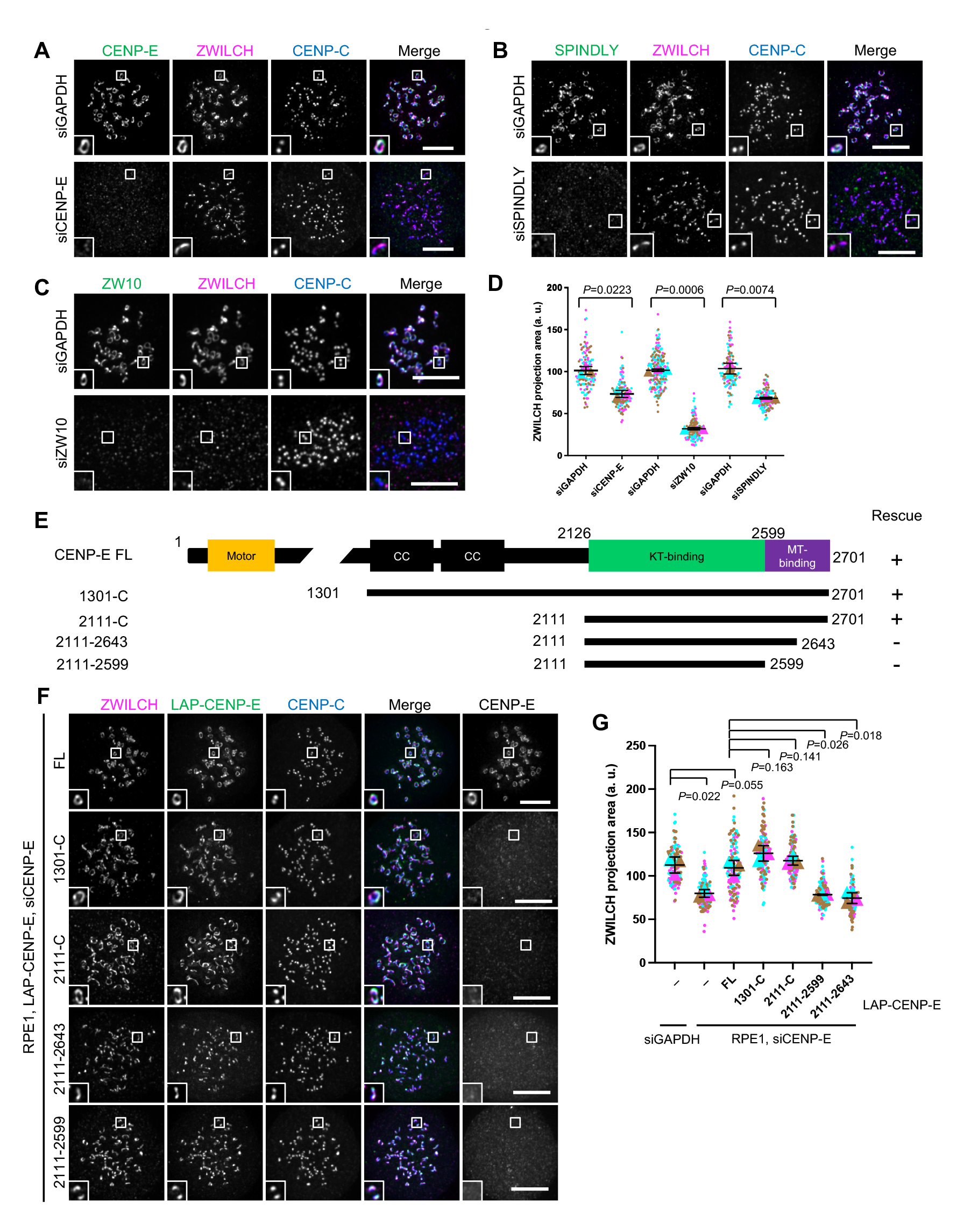
Fibrous corona expansion requires CENP-E. A: Immunostaining of ZWILCH (magenta), CENP-E (green) and CENP-C (blue) in control and CENP-E-depleted RPE1 cells treated with nocodazole overnight. Bar, 5 μm. B: Immunostaining of ZWILCH (magenta), SPINDLY (green) and CENP-C (blue) in control and SPINDLY-depleted RPE1 cells treated with nocodazole overnight. Bar, 5 μm. C: Immunostaining of ZWILCH (magenta), ZW10 (green) and CENP-C (blue) in control and ZW10-depleted RPE1 cells treated with nocodazole overnight. Bar, 5 μm. D: Projection area of ZWILCH in control, CENP-E-depleted, ZW10-depleted and SPINDLY-depleted RPE1 cells treated with nocodazole overnight. Student’s test. E: Domain organization of CENP-E and the deletion mutants used to rescue fibrous corona. CC, coiled coil. Rescue of fibrous corona are indicated. F: Immunostaining of ZWILCH (magenta), CENP-E (gray) and CENP-C (blue) in CENP-E-depleted RPE1 cells that over-express the indicated LAP-tagged CENP-E mutant after nocodazole treatment overnight. Bar, 5 μm. Rabbit polyclonal CENP-E antibody recognizes human CENP-E 955-1571. G: Projection area of ZWILCH in control and CENP-E-depleted RPE1 cells and CENP-E-depleted RPE1 cells that inducibly over-express the indicated LAP-tagged CENP-E mutant with nocodazole treatment overnight. Student’s test.

To determine how CENP-E contributes to fibrous corona formation, we next performed RNAi complementation with a series of LAP-tagged CENP-E variants (Figure 1E). This revealed that the motor domain of CENP-E was dispensable for fibrous corona formation (Figure 1F, G). Consistently, the fibrous corona was unaffected by treatment with GSK923295, a small molecular inhibitor of CENP-E motor activity (Figure S1G) (Qian et al., 2010). Instead, a fragment of CENP-E encompassing amino acids 2111-2701 (hereafter named 2111-C), including the kinetochore-binding (KT-binding) and microtubule-binding (MT-binding) domains, was sufficient to rescue fibrous corona formation in the absence of endogenous CENP-E. This depended on the C-terminal 58 amino acids within the MT-binding domain (Figure 1F, G, S1H, I).

### Fibrous corona formation requires C-terminal farnesylation of CENP-E

Having found that the C-terminal-most part of CENP-E (2111-C) was sufficient for fibrous corona formation, we next attempted to further narrow down the essential region. Expression in HeLa cells of a series of MT-binding domain mutants of CENP-E 2111-C (Figure 2A) revealed that only the C-terminal 12 amino acids were critical for fibrous corona formation (Figure 2B, C). This sequence contains a CAAX box, which is a farnesylation motif (Ashar et al., 2000). We have previously shown that cells treated with the farnesylation inhibitor lonafarnib do not expand fibrous coronas, a phenotype also seen in cells expressing a SPINDLY mutant that cannot be farnesylated (Sacristan et al., 2018). To examine a contribution of CENP-E farnesylation, we substituted the C-terminal 12 amino acids of CENP-E one by one for alanine (Figure 2D) and observed that only C2698A and Q2701A, the two residues of the CAAX box critical for farnesylation, did not support fibrous corona expansion in HeLa and RPE1 cells (Figure 2E, F, S2A, B, C). We conclude that farnesylation of CENP-E is required for fibrous corona formation.

**Figure 2.**
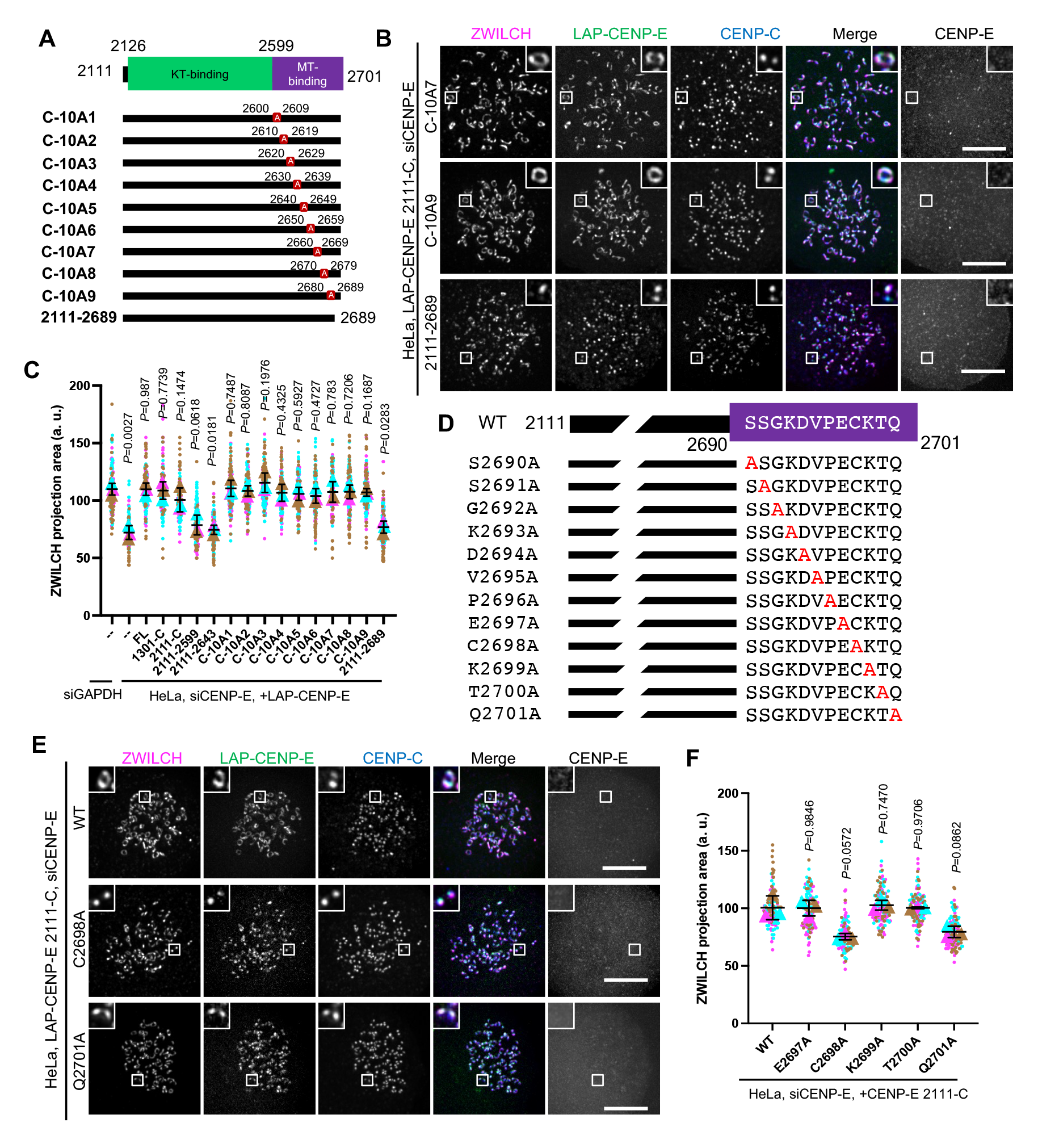
Fibrous corona formation requires C-terminal farnesylation of CENP-E. A: Domain organization of CENP-E 2111-C and various respective mutants. 2111-C fragment of CENP-E with different stretches of 10 amino acids were substituted for alanine residues or was deleted of the C-terminal 12 amino acids. B: Immunostaining of ZWILCH (magenta), CENP-E (gray) and CENPC (blue) in CENP-E-knockdown HeLa cells that over-express the indicated CENP-E mutant after 8hrs nocodazole treatment. Bar, 5 μm. Rabbit polyclonal CENP-E antibody recognizes human CENP-E 955- 1571. C: Projection area of ZWILCH in control and CENP-E-knockdown HeLa cells and CENP-E-knockdown HeLa cells that inducibly over-express the indicated LAP-tagged CENP-E mutant after 8hrs nocodazole treatment. Student’s test. D: Domain organization of CENP-E C-terminal and the indicated point-mutated mutants used to rescue fibrous corona. E: Immunostaining of ZWILCH (magenta), CENP-E (gray) and CENPC (blue) in CENP-E-knockdown HeLa cells that over-express the indicated LAP-tagged CENP-E mutant after 8hrs nocodazole treatment. Bar, 5 μm. Rabbit polyclonal CENP-E antibody recognizes human CENP-E 955-1571. F: Projection area of ZWILCH in CENP-E-knockdown HeLa cells that over-express the indicated LAP-tagged CENP-E mutant after 8hrs nocodazole treatment. Student’s test.

### Farnesylated CENP-E is essential for kinetochore-derived microtubule nucleation

Our prior work showed that fibrous coronas are the site of kinetochore-dependent microtubule nucleation, which aids spindle assembly and chromosome congression (Wu et al., 2023). To verify the requirement of CENP-E for fibrous corona expansion, we next assessed whether kinetochore-derived microtubule nucleation depends on CENP-E. To this end, we washed out nocodazole from RPE1 cells that had been treated with nocodazole overnight. In contrast to control cells, which had abundant microtubules newly nucleated on kinetochores 3 minutes after washout of nocodazole, cells depleted of CENP-E showed virtually no microtubule nucleation on kinetochores (Figure 3A, B, S3A). Consistent with our prior observation that fibrous corona-derived microtubule nucleation requires the LIC1 subunit of dynein (Wu et al., 2023), depletion of CENP-E substantially removed LIC1 from kinetochores (Figure S3B, C). Inhibition of CENP-E motor activity by treatment with GSK923295 did not prevent kinetochore-derived microtubule nucleation (Figure S3D, E), and, consistently, neither did expression of the 2111-C fragment, which lacks the motor domain (Figure 3C, D). These results indicate that CENP-E promotes microtubule nucleation at kinetochores through its non-motor role in fibrous corona expansion. In support of this, the mutants of CENP-E lacking the CAAX box (2111-2689) or in which the farnesyl-acceptor cysteine is substituted (C2698A) failed to restore microtubule nucleation (Figure 3C, D, S3F, G). Together, our findings show that CENP-E is essential for kinetochore-dependent microtubule nucleation by ensuring expanded fibrous coronas.

**Figure 3.**
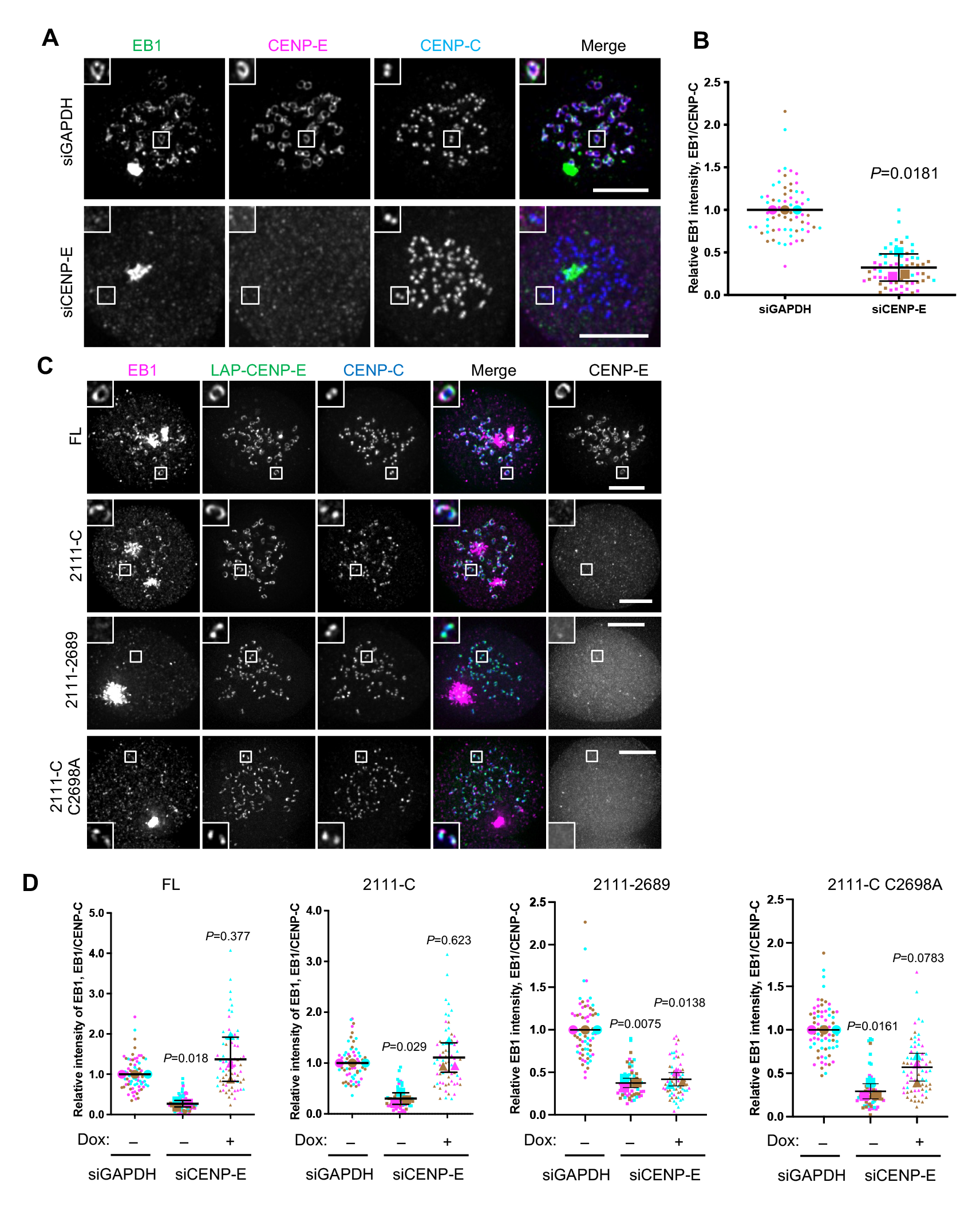
Farnesylated CENP-E is essential for kinetochore-derived microtubule nucleation. A: Immunostaining of EB1 (green), CENP-E (magenta) and CENPC (blue) in control and CENP-E-knockdown RPE1 cells after recovery of 3minutes from nocodazole washout. Bar, 5 μm. B: Relative EB1 intensity at the kinetochore compared to CENPC in control and CENP-E-knockdown RPE1 cells after recovery of 3minutes from nocodazole washout. Student’s test. C: Immunostaining of EB1 (magenta), CENP-E (gray) and CENPC (blue) in CENP-E-knockdown RPE1 cells that over-express the indicated CENP-E mutant after recovery of 2.5∼3 minutes from nocodazole washout. Bar, 5 μm. Rabbit polyclonal CENP-E antibody recognizes human CENP-E 955-1571. D: Relative EB1 intensity at the kinetochore compared to CENPC in control and CENP-E-knockdown RPE1 cells that inducibly over-express the indicated CENP-E mutant after recovery of 2.5∼3minutes from nocodazole washout. Student’s test.

### A farnesylated CENP-E fragment can recruit the RZZ-S complex in interphase cells

During our experiments, we noticed that the 2111-C variant of CENP-E and occasionally also longer variants (Figure S4A), which all include a microtubule-binding domain, could be found to colocalize with stable microtubules in interphase cells (Figure 4A). Surprisingly, we found that RZZ-S complex proteins localized to these 2111-C-covered microtubules (Figure 4B, S4A). Recruitment of RZZ-S (marked by ZWILCH) to these microtubules depended on farnesylation of the 2111-C fragment (Figure 4C, D, S4B). Cells treated with nocodazole to deplete microtubules still retained small clusters of 2111-C colocalizing with ZWILCH, in a manner depended on 2111-C farnesylation (Figure S4C). We thus conclude that farnesylated 2111-C is sufficient to recruit RZZ-S to ectopic sites in interphase cells. To further explore the mechanisms by which RZZ-S is recruited to ectopic 2111-C locations, we performed proximity biotinylation proteomics on interphase HeLa cells expressing 2111-C variants tagged with TurboID at their amino-termini (to not interfere with carboxy-terminal farnesylation) (Branon et al., 2018). Compared to the biotinylated fractions from cells expressing unfarnesylated 2111-C variants (2111-2689, C2698A), those from cells expressing farnesylated 2111-C variants (C-10A7, C-10A9) were consistently enriched for SPINDLY (SPDL1) (Figure 4E, S4D, E). Farnesylated 2111-C might thus, directly or indirectly, recruit RZZ-S complexes via SPINDLY.

**Figure 4.**
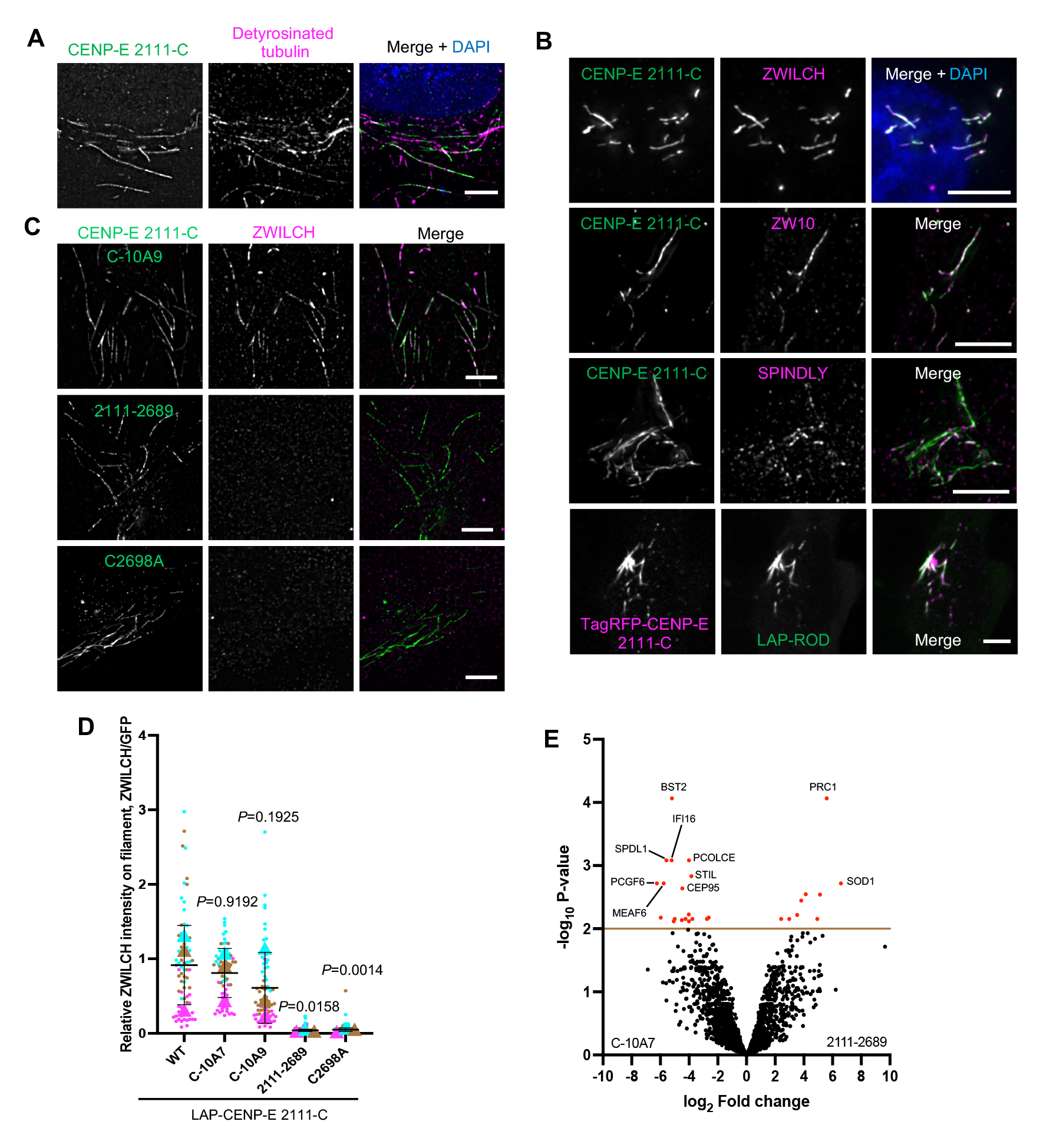
A farnesylated CENP-E fragment can recruit the RZZ-S complex in interphase cells. A: Immunostaining of detyrosinated tubulin (magenta) in RPE1 cells that over-express LAP-CENP-E 2111-C. Bar, 5 μm. B: Immunostaining of ZWILCH/ZW10/SPINDLY (magenta) in HeLa cells that over-express LAP-CENP-E 2111-C (top 3 panels); HeLa cells that co-express TagRFP-CENP-E 2111-C and LAP-ROD. Bar, 5 μm. C: Immunostaining of ZWILCH (magenta) in RPE1 cells that over-express the indicated LAP-tagged CENP-E mutants. Bar, 5 μm. D: Relative intensity of ZWILCH on the microtubule filaments bound with indicated CENP-E mutants. Student’s test. E: Volcano plot shows top enriched proteins bound to the streptavidin beads incubation with extracts of HeLa cells expressing LAP-TurboID-CENP-E 2111-C C-10A7 (left) or LAP-TurboID-CENP-E 2111-2689 (right).

### Farnesylation promotes interaction of endogenous CENP-E with fibrous corona proteins in mitosis

Having observed that over-expressed 2111-C recruits RZZ-S in interphase cells, we next wished to examine whether endogenous CENP-E interacts with RZZ-S in mitosis.

Kinetochores in cells with monopolar spindles (due to treatment with the Eg5 inhibitor STLC (Brier et al., 2004)) have small fibrous coronas (likely as a result of ongoing dynein-mediated poleward stripping) with detectable amounts of ZW10 and CENP-E (Figure 5A, C) (Famulski et al., 2011; Howell et al., 2001; Sacristan et al., 2018). Unexpectedly, additional inhibition of CENP-E with GSK923295 resulted in displacement of CENP-E and ZW10 from kinetochores and accumulation on the spindle pole (Figure 5A, B, C). This polar accumulation was not a result of dynein motor activity (see siZW10 below), but, since motor-inactive CENP-E tends to tightly bind microtubules (Wood et al., 2010), instead may result from transport of the inhibited CENP-E by poleward microtubule flux. Regardless of the exact mechanism, this phenotype provided us with a useful assay to examine potential kinetochore proteins that co-accumulate with CENP-E on spindle poles, and to examine the mechanism thereof. Besides CENP-E and ZW10, treatment with GSK923295 caused depletion from kinetochores and accumulation on the spindle pole of the fibrous corona proteins SPINDLY, ZWILCH, MAD1 and p150^Glued^, but not of the outer kinetochore proteins BUBRI and ZWINT-1 (Figure 5D, S5A). The kinetochore depletion and polar accumulation of ZW10 and ZWILCH, and therefore presumably of the other fibrous corona proteins, depended on CENP-E (Figure 5E, F, S5F, G). In contrast, ZW10 RNAi did not impact CENP-E accumulation on the spindle pole (Figure 5E, G). Furthermore, ZWILCH depletion prevented SPINDLY accumulation, and vice versa (Figure 5H, I, J), showing that RZZ-S members behaved as one complex. We conclude that in cells with monopolar spindles, transport of inactive CENP-E from kinetochores to spindle poles causes translocation of fibrous corona proteins. Importantly, the ability of inactive CENP-E to translocate RZZ-S was dependent on its C-terminal farnesylation: lonafarnib treatment of RPE1 cells (Figure 5K, L, S5B, C, D, E) or expression of unfarnesylated variants of CENP-E in CENP-E-depleted HeLa cells (Figure S5F, G) abolished translocation of RZZ-S and MAD1 to spindle poles.

**Figure 5.**
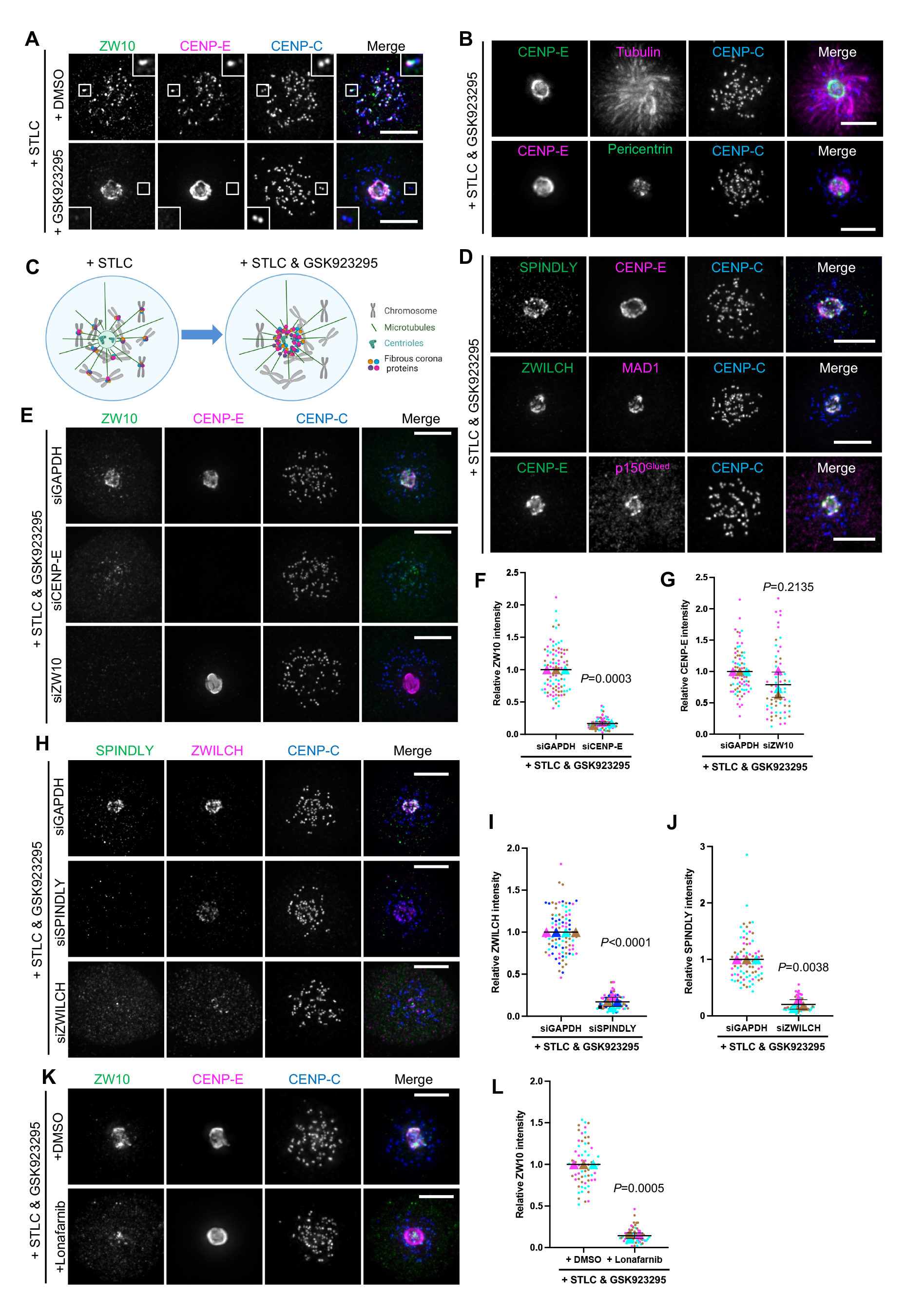
Farnesylation promotes interaction of endogenous CENP-E with fibrous corona proteins in mitosis. A: Immunostaining of ZW10 (green), CENP-E (magenta) and CENPC (blue) in STLC-treated RPE1 cells co-treated with DMSO or GSK923295 overnight. Bar, 5 μm. B: Immunostaining of CENP-E (green) / Pericentrin (green), Tubulin (magenta) / CENP-E (magenta) and CENPC (blue) in RPE1 cells treated with STLC and GSK923295 overnight. Bar, 5 μm. C: Cartoon showing the effects on fibrous corona proteins localization of indicated drugs treatment. D: Immunostaining of SPINDLY (green) / ZWILCH (green) / CENP-E (green), CENP-E (magenta) / MAD1 (magenta) / p150^Glued^ (magenta) and CENPC (blue) in RPE1 cells treated with STLC and GSK923295 overnight. Bar, 5 μm. E: Immunostaining of ZW10 (green), CENP-E (magenta) and CENPC (blue) in control, CENP-E-depleted, and ZW10-depleted RPE1 cells treated with STLC and GSK923295 overnight. Bar, 5 μm. F: Relative intensity of ZW10 at the spindle pole in control and CENP-E-depleted RPE1 cells treated with STLC and GSK923295 overnight. G: Relative intensity of CENP-E at the spindle pole in control and ZW10-depleted RPE1 cells treated with STLC and GSK923295 overnight. H: Immunostaining of SPINDLY (green), ZWILCH (magenta) and CENPC (blue) in control, SPINDLY-depleted, and ZWILCH-depleted RPE1 cells treated with STLC and GSK923295 overnight. Bar, 5 μm. I: Relative intensity of ZWILCH at the spindle pole in control and SPINDLY-depleted RPE1 cells treated with STLC and GSK923295 overnight. J: Relative intensity of SPINDLY at the spindle pole in control and ZWILCH-depleted RPE1 cells treated with STLC and GSK923295 overnight. K: Immunostaining of ZW10 (green), CENP-E (magenta) and CENPC (blue) in RPE1 cells treated with STLC and GSK923295 overnight in the presence of DMSO or 5 μM lonafarnib. Bar, 5 μm. L: Relative intensity of ZW10 at the spindle pole in RPE1 cells treated with STLC and GSK923295 overnight in the presence of DMSO or 5 μM lonafarnib.

### CENP-E impacts fibrous corona formation after initial RZZ-S kinetochore recruitment

Our data thus far showed that farnesylated CENP-E is important for fibrous corona formation and that CENP-E can quite strongly interact with RZZ-S and/or other fibrous corona components in a manner dependent on its farnesylation. To understand more about how CENP-E promotes fibrous corona formation, we asked when and how CENP-E localizes to kinetochores, relative to RZZ-S. Again we made use of the monopolar spindle assay, in which fibrous corona proteins are depleted from kinetochores and accumulate at the spindle pole upon chemical inhibition of CENP-E. Subsequent addition of nocodazole to these cells to deplete microtubules caused detectable relocalization of ZW10 back to the kinetochores within ∼1 minute, followed by CENP-E after ∼8 minutes (Figure 6A, B, S6A). Importantly, CENP-E depletion did not affect the initial recruitment of ZW10 under these conditions, but did impact the ability of ZW10 to expand to the recognizable crescents of the fibrous corona, and vice versa (Figure 6C, D, E, S6B, C). CENP-E and ZW10 thus initially localized to kinetochores independently but required each other for fibrous corona expansion thereafter. CENP-E interacts with BUBR1 (Chan et al., 1998; Ciossani et al., 2018; Legal et al., 2020; Yao et al., 2000), and indeed depletion of BUBR1 abrogated the initial reappearance of CENP-E at kinetochores after nocodazole addition, but not its localization to fibrous coronas at later stages (Figure 6F, G, S6D, E). Co-depletion of ZW10 and BUBR1 blocked the early recruitment of CENP-E to kinetochores and inhibited subsequent fibrous corona expansion (Figure 6H, I, S6F). Together, these data suggest that CENP-E is recruited to kinetochores independently of and after RZZ-S and subsequently promotes RZZ-S-mediated fibrous corona expansion.

**Figure 6.**
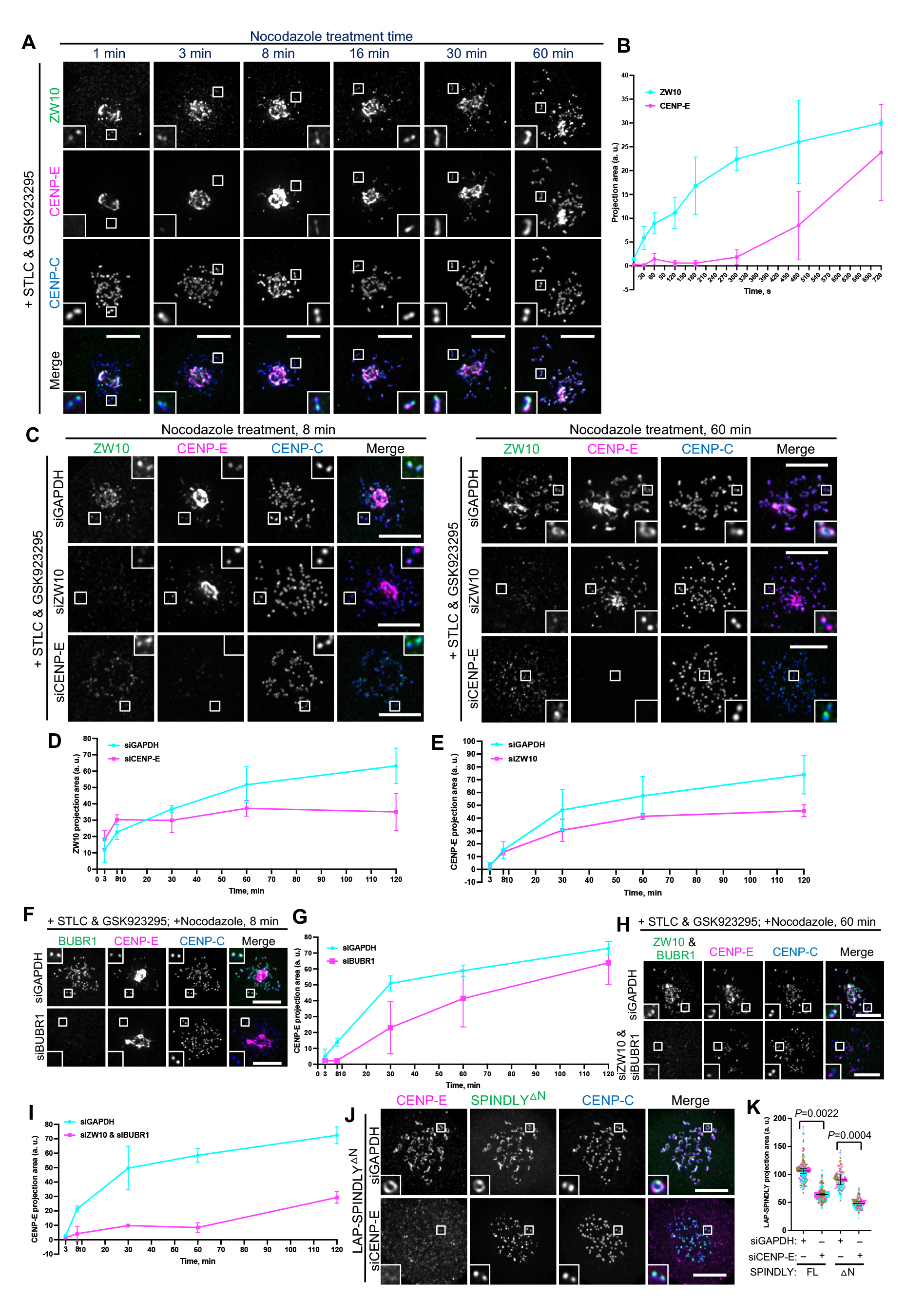
CENP-E impacts fibrous corona formation after initial RZZ-S kinetochore recruitment. A: Immunostaining of ZW10 (green), CENP-E (magenta) and CENP-C (blue) in RPE1 cells treated with nocodazole for indicated time, which had been arrested with STLC and GSK923295 overnight. Bar, 5 μm. B: Projection area of ZW10 and CENP-E in RPE1 cells treated with nocodazole for indicated time, which had been arrested with STLC and GSK923295 overnight. C: Immunostaining of ZW10 (green), CENP-E (magenta) and CENP-C (blue) in control, ZW10-depleted, and CENP-E-depleted RPE1 cells treated with nocodazole for indicated time, which had been arrested with STLC and GSK923295 overnight. Bar, 5 μm. D: Projection area of ZW10 in control and CENP-E-depleted RPE1 cells treated with nocodazole for indicated time, which had been arrested with STLC and GSK923295 overnight. E: Projection area of CENP-E in control and ZW10-depleted RPE1 cells treated with nocodazole for indicated time, which had been arrested with STLC and GSK923295 overnight. F: Immunostaining of BUBR1 (green), CENP-E (magenta) and CENP-C (blue) in control and BUBR1-depleted RPE1 cells treated with nocodazole for indicated time, which had been arrested with STLC and GSK923295 overnight. Bar, 5 μm. G: Projection area of CENP-E in control and BUBR1-depleted RPE1 cells treated with nocodazole for indicated time, which had been arrested with STLC and GSK923295 overnight. H: Immunostaining of ZW10 (green), BUBR1 (green), CENP-E (magenta) and CENP-C (blue) in control and ZW10 & BUBR1-co-depleted RPE1 cells treated with nocodazole for indicated time, which had been arrested with STLC and GSK923295 overnight. Bar, 5 μm. I: Projection area of CENP-E in control and ZW10 & BUBR1-co-depleted RPE1 cells treated with nocodazole for indicated time, which had been arrested with STLC and GSK923295 overnight. J: Immunostaining of CENP-E (magenta) and CENP-C (blue) in control and CENP-E-knockdown RPE1 cells that inducibly over-express LAP-SPINDLY Λ1N after nocodazole treatment overnight. Bar, 5 μm. K: Projection area of LAP-SPINDLY FL/Λ1N in control and CENP-E-knockdown RPE1 cells that inducibly over-express LAP-SPINDLY FL/Λ1N after nocodazole treatment overnight. Student’s test.

SPINDLY has a central role in RZZ-S oligomerization - which in vitro does not require CENP-E (Raisch et al., 2022; Sacristan et al., 2018) - that involves a conformational transition to relieve auto-inhibition (Sacristan et al., 2018). A conformationally ‘open’ SPINDLY (SPINDLY^ΔN^) bypasses the need for regulatory mechanisms of fibrous corona formation and does not bind dynein/dynactin, resulting in persistent fibrous coronas (Raisch et al., 2022; Sacristan et al., 2018). If CENP-E functions upstream of SPINDLY conformational ‘activation’, expression of SPINDLY^ΔN^ should restore fibrous coronas in CENP-E-depleted cells. As shown in Figure 6J, K and S6G, fibrous corona formation in SPINDLY^ΔN^-expressing cells still relied on CENP-E, showing CENP-E is not contributing to the mechanisms that control SPINDLY conformation. Given that CENP-E is also not needed for oligomerization per se (Raisch et al., 2022; Sacristan et al., 2018), these data suggest CENP-E impacts the fibrous corona in other ways, perhaps - since CENP-E is recruited later than RZZ-S - by stabilizing the RZZ-S meshwork during its formation.

### Comparative genomics of fibrous corona formation mechanisms in eukaryotes

To examine whether the mechanisms for fibrous corona formation might be conserved in eukaryotes, we built on our previous approach of comparative genomics of diverse eukarotic species (van Hooff et al., 2017). While ZW10 orthologs are omnipresent in eukaryotes, likely due to their primary role in vesicular trafficking (Sun et al., 2007), RZS (ROD/ZWILCH/SPINDLY) is an evolutionary coherent module in eukaryotes: there is a high degree of co-occurrence (presence or absence) of orthologs of its members in eukaryotic species, with co-presence of all RZZ-S complex members being restricted to Opisthokonta (fungi and animals) (van Hooff et al., 2017). We strengthened these previous observations in a larger, more diverse set of species genomes, utilizing refined homology detection methods (Figure S7). Given that RZZ-S form a biochemical unit in several animal model organisms, its coherent presence/absence profiles in eukaryotic species could signal the existence of a similar functional unit, and perhaps even of fibrous coronas, in the various species whose genomes encode RZZ-S orthologs. In contrast to RZS, CENP-E orthologs can be found in many species from all major eukaryotic lineages, similar to ZW10 (Figure 7) (van Hooff et al., 2017). However, CENP-E orthologous genes encoding a C-terminal CAAX box (defined by a cysteine at the −4 position from the C-terminus) appear only in two distantly-related groups: the Opisthokonta/Apusozoa, which unites fungi, animals and several unicellular relatives, and the Stramenopila/Alveolata (‘SA’) (Figure 7 and S7). Due to the large phylogenetic distance between these lineages, we infer that the acquisitions of CAAX box in CENP-E ancestors were likely independent. Importantly, SPINDLY is inferred to emerge at the base of Obazoa (Opisthokonta/Apusozoa/Breviata), and in this lineage CENP-E’s CAAX box has a high degree of co-occurrence with RZS orthologs (Figure 7 and S7). Emergence of the CAAX box in CENP-E near the base of the Obazoa therefore coincides with the emergence of SPINDLY and with the presence of the full RZZ-S unit.

**Figure 7.**
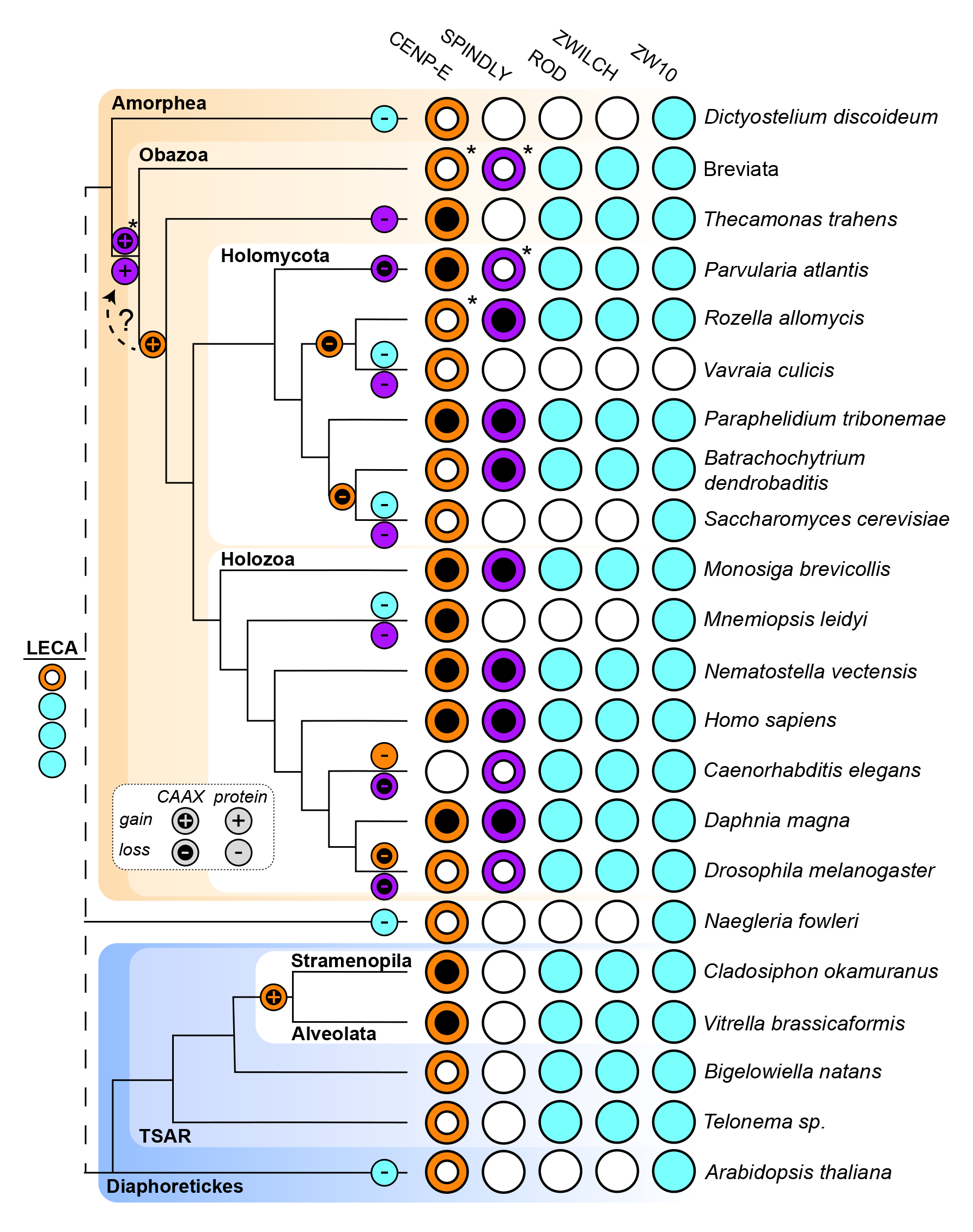
Comparative genomics of fibrous corona formation mechanisms in eukaryotes. Phylogenetic profiling of CENP-E, the CAAX box of CENP-E, SPINDLY, the CAAX box of SPINDLY, ROD, ZWILCH and ZW10 across eukaryotic diversity represented by selected species. Evolutionary reconstructions of loss and gain of proteins and CAAX boxes are projected onto the phylogenetic tree on the left-hand side of the figure. Presences are represented by colored circles and absences are represented by white circles. Due to the poor quality of the sequence data in the Breviata, the presence/absence data is the combination of inferences of two member species (*Pygsuia biforma* and *Lenisia limosa*). For inferring the presence of a CAAX box in either CENP-E or SPINDLY, having a complete protein sequence is essential. In cases where features of the predicted protein sequence strongly suggest that only a fragment is available, it is not possible to infer the presence of a CAAX box and such cases are indicated with an asterisk (*). These fragments affect in particular the inference of the exact timing of the CENP-E CAAX box, which emerged at the latest in the last common ancestor of Holozoa & Holomycota (combined the Opisthokonta) but may have already been present in the ancestor of the Breviata and the rest of the Obazoa, as is inferred for SPINDLY, as indicated with the dotted line with a question mark.

## Discussion

The fibrous corona is a meshwork that temporarily covers the outer kinetochores during early mitosis and meiosis and that assists in spindle assembly (Kops and Gassmann, 2020). Several recent studies have shown key mechanisms for how it expands and compacts. Central to these mechanisms is the RZZ-S complex, which can oligomerize into a filamentous meshwork in vitro (Pereira et al., 2018; Raisch et al., 2022; Sacristan et al., 2018). In cells, this oligomerization requires MPS1 activity and a conformational transition of SPINDLY, involving release of auto-inhibition imposed by N-terminal sequences (Raisch et al., 2022; Sacristan et al., 2018). We now show that fibrous corona formation in cells additionally requires the mitotic kinesin CENP-E. The contribution of CENP-E does not involve its motor activity, since both chemically inhibited CENP-E and truncated versions of CENP-E lacking the motor domain can substitute for endogenous CENP-E in supporting fibrous corona formation. Instead, its C-terminal ∼590 amino acids and farnesylation of cysteine 2698 suffice for this newly identified function.

Given that RZZ-S oligomerization is the driving mechanism for fibrous corona formation, it is likely that CENP-E’s contribution impinges on this process. Our proteomics analysis of CENP-E-interacting proteins and our observations that CENP-E recruits RZZ-S to ectopic sites in interphasic and mitotic cells suggests that CENP-E interacts with RZZ-S, possibly via SPINDLY. The CENP-E/RZZ-S interaction is most likely mediated by the KT-binding domain of CENP-E: it is included in the 2111-C fragment and is essential for binding of CENP-E to kinetochores and fibrous coronas (Chan et al., 1998). The interaction additionally requires farnesylation of CENP-E and this is also critical for fibrous corona formation. It is noteworthy that SPINDLY^ι1N^ partially rescues fibrous corona expansion in the absence of global farnesylation (Sacristan et al., 2018), but not in the absence of CENP-E (Fig. 6J). This suggests that CENP-E’s role in fibrous corona expansion is not solely mediated by CENP-E farnesylation, and may include additional interactions with RZZ-S. How does CENP-E achieve this? It could affect the regulation of RZZ-S oligomerization by MPS1, involving release of the auto-inhibition of SPINDLY (Raisch et al., 2022; Sacristan et al., 2018). However, if this were true, persistently uninhibited SPINDLY (SPINDLY^ι1N^) should be able to bypass this step in the absence of CENP-E. This is not so, showing CENP-E contributes to a parallel or downstream event. An alternative hypothesis is that CENP-E is an integral component of the RZZ-S meshwork. However, RZZ-S can oligomerize in vitro without CENP-E (Raisch et al., 2022; Sacristan et al., 2018). In kinetochores, however, the molecular environment is different and CENP-E might be required to promote the reaction, for example as a catalyst, although there is currently no clear molecular rationale for this. Based on our observation that CENP-E follows RZZ to kinetochores, perhaps a more likely model is one in which CENP-E stabilizes the RZZ-S meshwork during its expansion. SPINDLY interacts with RZZ via its farnesyl moiety that docks into a hydrophobic pocket in the ROD beta-propeller (Mosalaganti et al., 2017; Raisch et al., 2022). A similar mode of interaction could apply to farnesylated CENP-E. This could be through the known farnesyl binding pocket on ROD or a different region in the RZZ-S complex, for example SPINLDY, which was the RZZ-S member most proximal to farnesylated 2111-C in our proximity biotinylation experiments. Extensive biochemical reconstitution experiments with RZZ-S and CENP-E under different conditions are needed to reveal the exact molecular mechanism.

How conserved is the CENP-E contribution we identify in the present study? The fibrous corona structure as defined by electron microscopy imaging of unattached kinetochores has been observed in cells of various vertebrates (Bielek, 1978; Jokelainen, 1967; Rieder, 1982; Roos, 1973), but has not been examined as such in fungi and/or other non-animal lineages. Our deep homology searches for homologs of RZZ-S members revealed that the full complex emerges at the base of the Obazoa, with the emergence of ancestral SPINDLY. The core ingredients for fibrous corona formation were thus present in the ancestor to Obazoa. Although CENP-E orthologs are widespread in eukaryotes, the appearance of a farnesyl-acceptor cysteine residue at position −4 from the carboxy terminus in the Obazoa lineage shows a striking correlation with the emergence of SPINDLY (and its CAAX motif) and thus the full RZZ-S complex. Our current analysis pinpoints emergence of SPINDLY one branch earlier than the cysteine in CENP-E, but note that the CENP-E ortholog sequences in Breviata are incomplete. Better sampling of genomes from Breviata species might therefore reveal full-length CENP-E ORF sequences with a cysteine at −4 from the carboxy terminus, which would indicate near-simultaneous emergence of SPINDLY and the potential for CENP-E farnesylation. Beyond SPINDLY, the RZZ-S complex and the potential for CENP-E farnesylation show strongly cohesive presence/absence in Obazoa, with notable coherent losses close in the species tree. Interestingly, kinetochores in colchicine-treated *Drosophila* S2 cells do not appear to form clear fibrous coronas (Maiato et al., 2006), and their CENP-E and SPINDLY orthologs lack CAAX boxes. Our analyses revealed emergence of a cysteine at −4 in CENP-E also in the Stramenopiles/Alveolate lineage of eukaryotes. Given the simple trajectory to acquiring a single amino acid change and given the absence of the cysteine in all other lineages except Obazoa, it was likely acquired independently in the Stramenopiles/Alveolate lineage. Species in this lineage also have RZZ orthologs, raising the question of whether they may have the capacity to form a structure analogous to a fibrous corona, despite the apparent absence of a SPINDLY ortholog. Future cell biological and biochemical studies on these and other species will be important to understand conservation and diversity in fibrous corona form and function among eukaryotes.

## Materials and methods

### Cell culture

RPE1 FlpIn cells, HEK293T and HeLa FlpIn cells were cultured in Dulbecco’s Modified Eagle Medium/Nutrient Mixture F-12 (DMEM/F-12 GlutaMAX; Gibco 10565018) supplemented with 9% tetracycline-free fetal bovine serum and 100 μg /ml penicillin-streptomycin (Sigma P0781).

### Plasmids and cloning

LAP-SPINDLY FL and LAP-SPINDLY^△N^ were used previously (Sacristan et al., 2018). LAP-CENP-E full length and variants in pCDNA5 vector were cloned by PCR-based strategy. Variant C-10A1: CENP-E 2111-C with amino acids from 2600 to 2609 substituted for 10 alanine residues; variant C-10A2: CENP-E 2111-C with amino acids from 2610 to 2619 substituted for 10 alanine residues; variant C-10A3: CENP-E 2111-C with amino acids from 2620 to 2629 substituted for 10 alanine residues; variant C-10A4: CENP-E 2111-C with amino acids from 2630 to 2639 substituted for 10 alanine residues; variant C-10A5: CENP-E 2111-C with amino acids from 2640 to 2649 substituted for 10 alanine residues; variant C- 10A6: CENP-E 2111-C with amino acids from 2650 to 2659 substituted for 10 alanine residues; variant C-10A7: CENP-E 2111-C with amino acids from 2660 to 2669 substituted for 10 alanine residues; variant C-10A8: CENP-E 2111-C with amino acids from 2670 to 2679 substituted for 10 alanine residues; variant C-10A9: CENP-E 2111-C with amino acids from 2680 to 2689 substituted for 10 alanine residues; variant 2111-2689: CENP-E 2111-2689.

LAP-tagged CENP-E 1-426^2111-C variants were generated by fusing the N-terminal CENP-E 1-426 (comprising the motor domain) to the C-terminal CENP-E 2111-C (comprising KT-binding domain and MT-binding domain).

LAP-ROD in pCDNA5 vector was cloned based on the ROD cDNA, which was a kind gift from Reto Gassmann (Instituto de Biologia Molecular e Celular, Portugal), TagRFP-CENP-E 2111-C were cloned into the pLVX-IRES-Puro vector (Clontech) by PCR-based Gibson assembly method.

### Virus production

Lentiviruses were produced by co-transfection of HEK 293T cells with the lentiviral vector containing TagRFP-CENP-E 2111-C and separate plasmids that express Tat, VSV-G, Rev and Gag-Pol, with the transfection reagent of Fugene HD (Promega, E2311). 72 hours after transfection, cell culture supernatant was harvested and filtered.

### Stable cell lines and generation

RPE1 FlpIn cell line stably expressing EB3-TagRFPT-ires-GFP-ZWILCH was used previously (Wu et al., 2023).

RPE1 FlpIn cell line inducible expressing LAP-SPINDLY FL and LAP-SPINDLY^△N^ were used previously (Wu et al., 2023).

RPE1 FlpIn cell line were co-transfected with pCDNA5-constructs with gene of interest and construct of pOG44 recombinase in a 1:2 ratio with electroporation machine (Amaxa™ Nucleofector™ II) using protocol U-017. 3 days after transfection, RPE1 cells were selected with 100 μM hygromycin (Roche, 10843555001) for about 3 weeks.

HeLa FlpIn were co-transfected with pCDNA5 constructs with gene of interest and construct of pOG44 recombinase in a 1:9 ratio with Fugene HD (Promega, E2311) under the manufacturer’s instructions. One day after transfection, HeLa cells were selected with 200 μM hygromycin (Roche, 10843555001) for about 3 weeks.

To generate HeLa FlpIn cell stably expressing TagRFP-CENP-E 2111-C and inducible expressing LAP-ROD, HeLa FlpIn cells inducible expressing LAP-ROD were infected with lentiviruses and selected with 1 μg/mL puromycin for 1 week.

### siRNAs and transfection

The siRNAs targeting GAPDH (Dharmacon, D-001830-01-05), CENP-E (5’- CCACUAGAGUUGAAAGAUA-3’)(Kim et al., 2010), ZW10 (5’-UGAUCAAUGUGCUGUUCAA-3’)(Sacristan et al., 2018), Spindly (5’-GAAAGGGUCUCAAACUGAA-3’)(Sacristan et al., 2018), Zwilch (5’- UCUACAACGUGGUGAUAUA-3’)(Sacristan et al., 2018), BUBR1 (5′-AGAUCCUGGCUAACUGUUC −3′)(Smith et al., 2019) were purchased from Dharmacon. RPE1 cells or HeLa FlpIn cells were transfected with siRNAs at 100 nM using the HiPerFect (Qiagen), according to manufacturer’s instructions.

### Drug treatments on cells

RPE1 cells or HeLa FlpIn cells were treated with 6.6 μM nocodazole (Sigma-Aldrich, M1404) for 15hrs (overnight) for fibrous corona formation before the fixation. For cells transfected with siRNA, nocodazole was added 48hrs after siRNA transfection. For the rescue experiments, 1 μg/ml doxycycline (Sigma-Aldrich, D9891) was added to cells 40hrs after siRNA transfection to induce the expression of protein variants. For nocodazole washout assay, 6.6 μM nocodazole was added to the culture medium for 15hrs before washout. For CENP-E inhibition, cells were treated with 250nM GSK923295 (Selleck Chem, S7090) for 15hrs (overnight). For farnesyl transferase inhibition, cells were treated with 5uM lonafarnib (Selleckchem, S2797) for 15hrs (overnight). For the fast disassembly of microtubules, cells were treated with 13.2 μM nocodazole for 4 hours. Cells were treated with 10 mM STLC (Tocris Bioscience 1291) for 15hrs (overnight). For the experiments in which translocated fibrous corona proteins were released from spindle poles, cells were co-treated with STLC and GSK923295 for 15hrs (overnight), and were further treated with 13.2 μM nocodazole for indicated time before fixation. In the TurboID pulldown experiment, HeLa cells were treated with 1 μg/ml doxycycline for 24hrs and one hour of 250 μM biotin (Sigma-Aldrich, B4501-5G) before harvest.

### Antibodies

We used Guinea pig polyclonal antibody against CENPC (MBL, PD030), and rabbit polyclonal antibodies against CENP-E (Brown et al., 1996), ZW10 (Abcam, ab21582), SPINDLY (Bethyl, A301-354A-1), ZWILCH (a gift from Andrea Musacchio, Max Planck Institute of Molecular Physiology), detyrosinated α -tubulin (Abcam, ab48389), BUBR1 (Bethyl, A300-995A), pericentrin (Abcam, ab4448), MAD2 (custom, B. O. 017), ZWINT-1 (Abcam, ab71982). We used mouse monoclonal antibodies against α-tubulin (Sigma-Aldrich, T9026), MAD1 (Merck Millipore, MABE867), EB1 (BD Biosciences, 610534), CENPE (Abcam, ab5093), ZWILCH (a gift from Andrea Musacchio, Max Planck Institute of Molecular Physiology), p150^Glued^ (BD Biosciences, 612708). We used the following secondary antibodies: Goat anti-guinea pig Alexa Fluor 647 (Invitrogen, A21450), Goat anti-rabbit Alexa Fluor 488 (Invitrogen, A11034), Goat anti-rabbit Alexa Fluor 568 (Invitrogen, A11036), Goat anti-rabbit Alexa Fluor 405 (Invitrogen, A-31556), Goat anti-mouse Alexa Fluor 568 (Invitrogen, A11031), Goat anti-mouse Alexa Fluor 647 (Invitrogen, A21236) and GFP-Booster Atto 488 (Chromotek, gba-488).

### Immunofluorescence staining

Immunofluorescence method was described previously (Wu et al., 2023). In brief, cells grown on 12 mm coverslips were fixed with −20 °C methanol. After fixation, cells on coverslips were washed 3 times with PBS, followed by permeabilization with 0.1% triton X- 100 in PBS for 2 minutes, then followed by washing 3 times in PBS supplemented with 0.05% Tween 20 (PBST), and were blocked with 2% BSA diluted in PBST for 30 mins.

After blocking, cells were incubated with primary antibodies diluted in PBST containing 2% BSA for one hour at room temperature in humid condition. Subsequently, cells were washed three times with PBST, and were incubated with secondary antibodies diluted in PBST together with or without DAPI for 1hour at room temperature. Then, cells were washed three times with PBST. In the end, coverslips were sequentially rinsed in 70% and 100% ethanol, air-dried and mounted on glass slides with Prolong Gold antifade.

### Image acquisition and quantification

All images were acquired on a deconvolution system (DeltaVision Elite Applied Precision / GE Healthcare) with a 100X /1.40 NA UPlanSApo objective (Olympus) using SoftWorx 6.0 software (Applied Precision/GE Healthcare). Images were acquired as z-stacks at 0.2 μm intervals for 32 stacks and deconvolved using SoftWoRx. The images are maximum intensity projections of deconvoluted stacks.

Images were analysed with Fiji (https://fiji.sc). For quantification of proteins levels, all images of immunostaining experiments were acquired with identical illumination settings. Protein levels near the kinetochores were determined on maximum projections of z-stacks images using an ImageJ macro (Saurin et al., 2011; Wu et al., 2023) which thresholds the CENPC signal within the DAPI area or within the cytoplasm area in the images without DAPI staining. For quantification of EB1 levels on the kinetochores, the centrosome area was excluded as previous work (Wu et al., 2023). For fibrous corona size measurement, we determined Zwilch, MAD1, CENPE or ZW10 maximum projection area by an ImageJ macro that was used previously (Wu et al., 2023).

### Live-cell imaging

Imaging of RPE1 cells co-expressing EB3-TagRFPT and GFP-Zwilch in nocodazole washout assay was described previously (Wu et al., 2023). In brief, RPE1 cells were cultured in 24- well glass bottom plate (Cellvis, P24-0-N). Live-cell imaging was performed by acquiring images every 20 or 30 seconds at 1 × 1 binning on an Andor CSU-W1 spinning disk (50 μm disk) with Nikon 100× 1.45 NA oil objective. 488 nm and 561 nm lasers were used for sample excitation and images were acquired using an Andor iXon-888 EMCCD camera. Nocodazole washout was done during the imaging with warm culture medium.

### TurboID-based proteins pulldown for proteomics

In the TurboID-based proteins pulldown experiments, HeLa cells inducible expressing LAP-TurboID-CENP-E 2111-C variants were treated with 1 μg/mL doxycycline for 24hrs to induce the expression of LAP-TurboID-CENP-E 2111-C variants, and one hour of 250 μM biotin before harvest. Harvest the cells by trypsinization and centrifuge. Wash the cells once with ice-cold PBS, and lysate the cells with lysis buffer (50 mM HEPES pH 7.4, 150 mM NaCl, protease inhibitor, phosphatase inhibitor, 0.5 % Triton X-100) on ice for 10 min. After lysis, the samples were centrifuged at 14000rpm for 20 min at 4°C. The supernatants of the centrifuged samples were incubated with the streptavidin magnetic beads (Thermo Scientific, 88816) for 1hr in 4°C cold room. Wash the beads with wash buffer (50 mM HEPES pH 7.4, 150 mM NaCl, 0.1 % Triton X-100) 3 times. Remove the wash buffer after final wash, and store the beads in −80°C freezer.

### Liquid chromatography–mass spectrometry/ mass spectrometry

Precipitated proteins were denatured and alkylated in 50 µl 8 M Urea, 1 M ammonium bicarbonate (ABC) containing 10 mM TCEP (tris (2-carboxyethyl) phosphine hydrochloride) and 40 mM 2-chloro-acetamide for 30 minutes. After 4-fold dilution with 1 M ABC and digestion with trypsin (20 µg/200 µl), peptides were separated from the beads and desalted with homemade C-18 stage tips (3 M, St Paul, MN), eluted with 80% Acetonitrile (ACN) and, after evaporation of the solvent in the speedvac, redissolved in buffer A (0,1% formic acid). After separation on a 30 cm pico-tip column (75 µm ID, New Objective) in-house packed with C-18 material (1.9 µm aquapur gold, dr. Maisch) using a 140 minutes gradient (7% to 80% ACN, 0.1% FA), delivered by an easy-nLC 1000 (Thermo), peptides were electro-sprayed directly into a Orbitrap Fusion Tribrid Mass Spectrometer (Thermo Scientific). The latter was set in data dependent Top speed mode with a cycle time of 1 second, in which the full scan over the 400-1500 mass range was performed at a resolution of 240000. Most intense ions (intensity threshold of 15000 ions) were isolated by the quadrupole where after they were fragmented with a HCD collision energy of 30%. The maximum injection time of the ion trap was set to 50 milliseconds with injection of ions for all available parallelizable time. Raw data was analyzed with MaxQuant [version 1.6.3.4], using the Homo Sapiens (taxonomy ID: 9606) fasta file, extracted from UniprotKB. To determine proteins of interest, the protein groups output file was used to perform a differential enrichment analysis. Proteins with less than one unique peptide and proteins that have not been identified in at least two out of three of the replicates of one condition, were filtered out. Then, a background correction and normalization of the data was performed by variance stabilizing transformation; shifting and scaling the proteins intensities by sample group. A left-shifted Gaussian distribution randomization was used to impute since the data presented a pattern of missingness not at random (MNAR). Finally, a differential enrichment analysis was performed to identify those proteins that were differentially-enriched and selected those falling inside the threshold for log2 Fold Change and -log10 P-value higher than 2. The program used for the analyses was R [version 4.0.4] through RStudio [version 1.5.64].

The mass spectrometry proteomics data have been deposited to the ProteomeXchange Consortium via the PRIDE (Perez-Riverol et al., 2022) partner repository with the dataset identifier PXD040338.

### Eukaryotic sequence database

To profile the occurrence of orthologs of fibrous corona components across eukaryotes, we collected the predicted proteomes of 177 eukaryotic species representing the breath of currently-known eukaryotic diversity (External dataset 1). These 177 representative species were selected on several criteria: i) the overall quality of their predicted proteomes (as measured by BUSCO completeness (Simão et al., 2015)) (1), ii) their phylogenetic position in the eukaryotic tree of life, to ensure an as large as possible diversity within the dataset capturing all currently-defined eukaryotic supergroups and major evolutionary transitions, and iii) the overall interest of the biological community for certain (model) organisms, to ensure relevance and potentially future experimental verification of our inferences. These 177 species were selected from a larger set that combined several previously published eukaryotic proteome selections into a single set. These sets were taken from (Deutekom et al., 2019), the comparative set from EukProt (Richter et al., 2022), and the species selection of (Grau-Bové et al., 2022). The resulting dataset comprised 367 species. To select species from these 367, first all of them were subjected to the following analyses.

As typically splice variants are reported as separate sequences, we filtered splice variants out from the predicted proteomes and translated transcriptomes to produce non-redundant proteomes. Shortly, highly similar sequences from proteomes were clustered using the easy-cluster workflow from the MMseqs2 package v13.54111 (Steinegger and Söding, 2017). To evaluate whether the clusters strictly comprised true splice variants, we calculate the percentage identity across the alignment per cluster. First, sequences were aligned using MAFFT v7.505 (Katoh and Standley, 2013). Then, gap-rich regions were excluded from the alignment using the gappyout option of trimAl v1.4 (Capella-Gutiérrez et al., 2009), as these are possibly the product of in-or excluded exons resulting from alternative splicing. Finally, the percentage identity across all cluster members was calculated from the trimmed alignment with the sident option of trimAl v1.4 (Capella-Gutiérrez et al., 2009). Clusters containing sequences that all have at least 99% sequence identity to the longest transcript in the trimmed alignments were accepted and the longest transcript was taken as representative. If the percentage identity of a given sequence was below 99% to the cluster representative, the clustering was deemed spurious and the outlier sequences were considered as non-redundant sequences.

BUSCO (v5.2.2; eukarya_odb10) was used to determine the quality of the non-redundant proteomes of the selected species (Simão et al., 2015). Single-copy BUSCO orthologs found in at least 75% of all species were selected and taken as marker genes for the construction of a phylogenetic tree, yielding a comprehensive resolution of the diversity in this combined set. The phylogenetic tree was inferred using IQTree (v2.2.0) using the built-in SH-aLRT test and ModelFinder (Guindon et al., 2010; Kalyaanamoorthy et al., 2017; Minh et al., 2020), where the LG+F+R15 substitution model was selected. The resulting phylogeny was visualised in iTOL (Letunic and Bork, 2021) with BUSCO scores plotted on the leaves.

This herewith obtained representation of the phylogenetic relations and genome quality allowed for manual selection of species at key phylogenetic positions with the highest available quality proteomes across the relevant taxa. With a total of 177 species, the final set is computationally tractable while also allowing for maximum diversity and relevance to understand the evolution of proteins across eukaryotic lineages, such as the components of the fibrous corona.

### Homology detection and orthology assessment

Orthologs of CENP-E, SPINDLY, ROD, ZWILCH and ZW10 were detected according to previously described workflows (Tromer et al., 2019; van Hooff et al., 2017; van Hooff et al., 2019). Briefly, iterative profile hidden Markov model-based (HMM) homology searches were performed on the eukaryotic sequence database using the ‘hmmsearch’ method from the HMMER3 package v3.1b2 (Mistry et al., 2013). Here, we initially made use of previously-published HMMs by our group (Tromer et al., 2019). Homologous sequences were aligned using MAFFT E-INS-I v.7.505 (Katoh and Standley, 2013), alignments were trimmed using trimAl v1.4 to only include alignment positions with at least 10% occupancy to remove gap-rich and poorly-aligned regions (Capella-Gutiérrez et al., 2009), and finally, phylogenetic trees were inferred with FastTree 2 and IQ-Tree v2.2.0 to allow for phylogenomic assessment of homologous sequences (Minh et al., 2020; Price et al., 2010). In addition, we annotated putative orthologs using various homology-based methods including eggNOG-mapper v2 (Cantalapiedra et al., 2021), HHpred (Zimmermann et al., 2018), and BLASTP (Altschul et al., 1990). Besides detection of orthologous sequences in our current eukaryotic sequence database as described, we also included previously determined orthologs of fibrous corona components in Obazoan species not included in our initial selection of species to further increase the phylogenomic resolution in this clade based on (van Hooff et al., 2017).

Combined, our methods allow for the establishment of fine-grained manually-curated orthologous groups of fibrous corona components from our extended eukaryotic sequence database spanning the entire diversity of currently-known eukaryotic life on Earth. Alignments and HMMs of the orthologous groups as defined here are available via supplementary dataset.

### Gene (re)prediction

As gene prediction is imperfect, we scrutinised absences or incomplete sequences inferred from previously published predicted proteomes using the workflow described previously by (Tromer, 2017). Briefly, we interrogate genome assemblies for regions homologous to a missing ortholog using TBLASTN (Altschul et al., 1990). Here, we use previously-determined orthologs of closely-related species as query. If this method identified a homologous region in the genome, we performed gene prediction on this region using AUGUSTUS (Stanke and Morgenstern, 2005) and GENSCAN (Burge and Karlin, 1997) software using flexible settings to retrieve a set of possible transcripts which were then assessed manually against ortholog alignments to determine whether this represents a previously not predicted ortholog of this gene in this species. (Re)predicted sequences of orthologs are indicated as such in supplementary dataset.

## Acknowledgments

We thank R. Gassmann (Instituto de Biologia Molecular e Celular, Portugal), and A. Musacchio (Max Planck Institute of Molecular Physiology, Germany) for sharing reagents. We thank A. Musacchio (Max Planck Institute of Molecular Physiology, Germany), J. Welburn (University of Edinburgh, UK), M. Barisic (University of Copenhagen, Denmark) and the members of the Kops lab for discussions. The cartoon in figure 5C was created with biorender.com. The Kops lab is a member of the Oncode Institute, which is partly financed by the Dutch Cancer Society. This study was financially supported by grants from the European Research Council (ERC-SyG 855158) and the Dutch Cancer Society (11080). E. C. Tromer is supported by a personal fellowship from the Netherlands Organisation for Scientific Research (NWO VI. Veni. 202.223).

## Author contributions

J.W. and G.J.P.L.K. conceived the project. J.W. and G.J.P.L.K. designed all experiments, which were performed and analyzed by J.W., except for the proteomics experiments, which were performed and analyzed by P.S.A. and H.R.V.. M.R. performed comparative genomics, supervised by E.C.T. and B.S.. J.W. and G.J.P.L.K. wrote the manuscript, with input from all authors.

## Supplemental figures and legends

**Figure S1.**
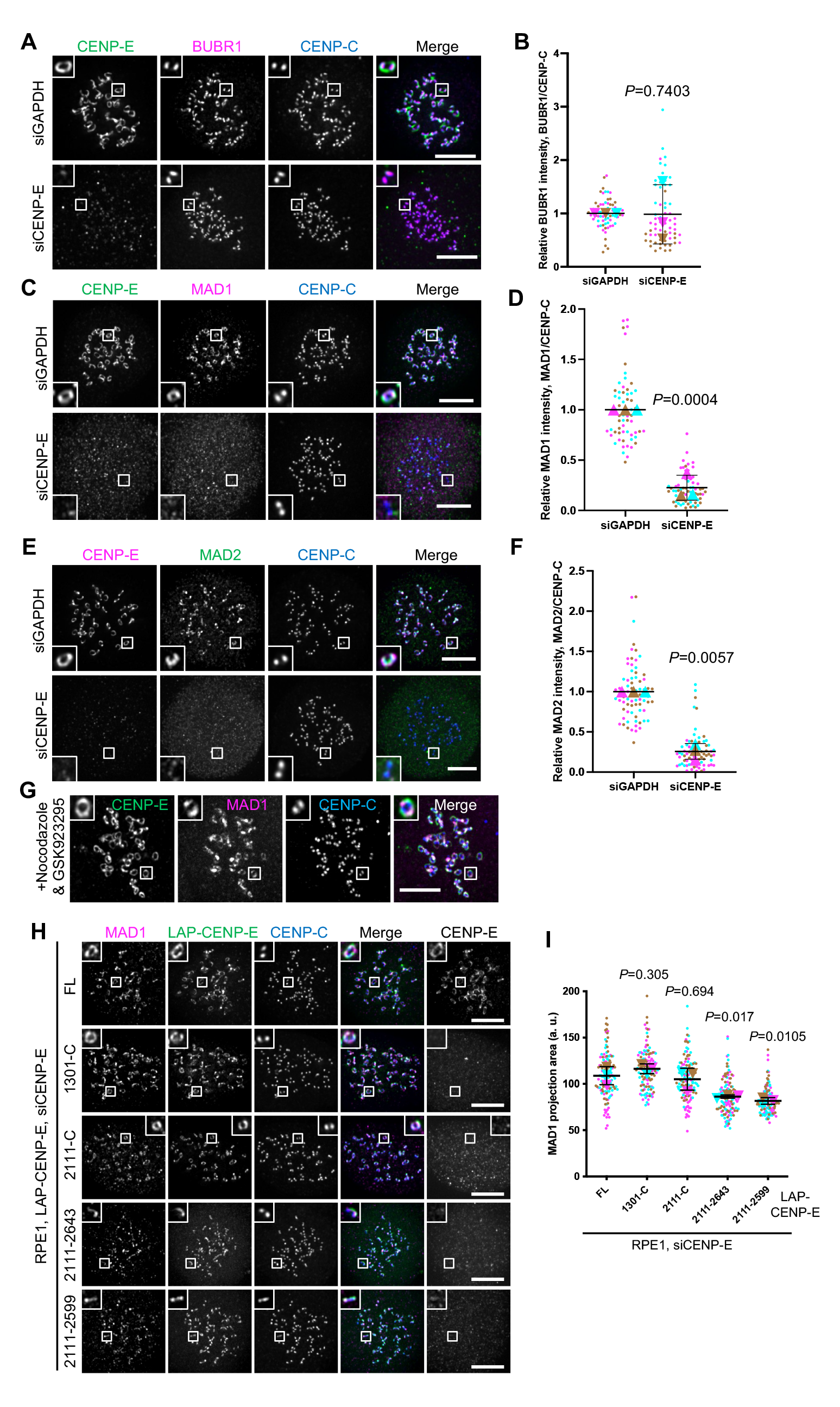
Fibrous corona expansion requires CENP-E. A: Immunostaining of BUBR1 (magenta), CENP-E (green) and CENPC (blue) in control and CENP-E-knockdown RPE1 cells treated with nocodazole overnight. Bar, 5 μm. B: Relative BUBR1 intensity at the kinetochore compared to CENPC in control and CENP-E-knockdown RPE1 cells treated with nocodazole overnight. Student’s test. C: Immunostaining of MAD1 (magenta), CENP-E (green) and CENPC (blue) in control and CENP-E-depleted RPE1 cells treated with nocodazole overnight. Bar, 5 μm. D: Relative intensity of MAD1 on the kinetochore in control and CENP-E-depleted RPE1 cells with nocodazole treatment overnight. Student’s test. E: Immunostaining of CENP-E (magenta), MAD2 (green) and CENPC (blue) in control and CENP-E-depleted RPE1 cells treated with nocodazole overnight. Bar, 5 μm. F: Relative intensity of MAD2 on the kinetochore in control and CENP-E-depleted RPE1 cells with nocodazole treatment overnight. Student’s test. G: Immunostaining of CENP-E (magenta), MAD1 (green) and CENPC (blue) in RPE1 cells co-treated with nocodazole and GSK923295 (inhibitor of CENP-E motor activity) overnight. Bar, 5 μm. H: Immunostaining of MAD1 (magenta), CENP-E (gray) and CENPC (blue) in CENP-E-depleted RPE1 cells that over-express the indicated LAP-tagged CENP-E mutant after nocodazole treatment overnight. Bar, 5 μm. Rabbit polyclonal CENP-E antibody recognizes human CENP-E 955-1571. I: Projection area of MAD1 in CENP-E-depleted RPE1 cells that inducibly over-express the indicated LAP-tagged CENP-E mutant with nocodazole treatment overnight. Student’s test.

**Figure S2.**
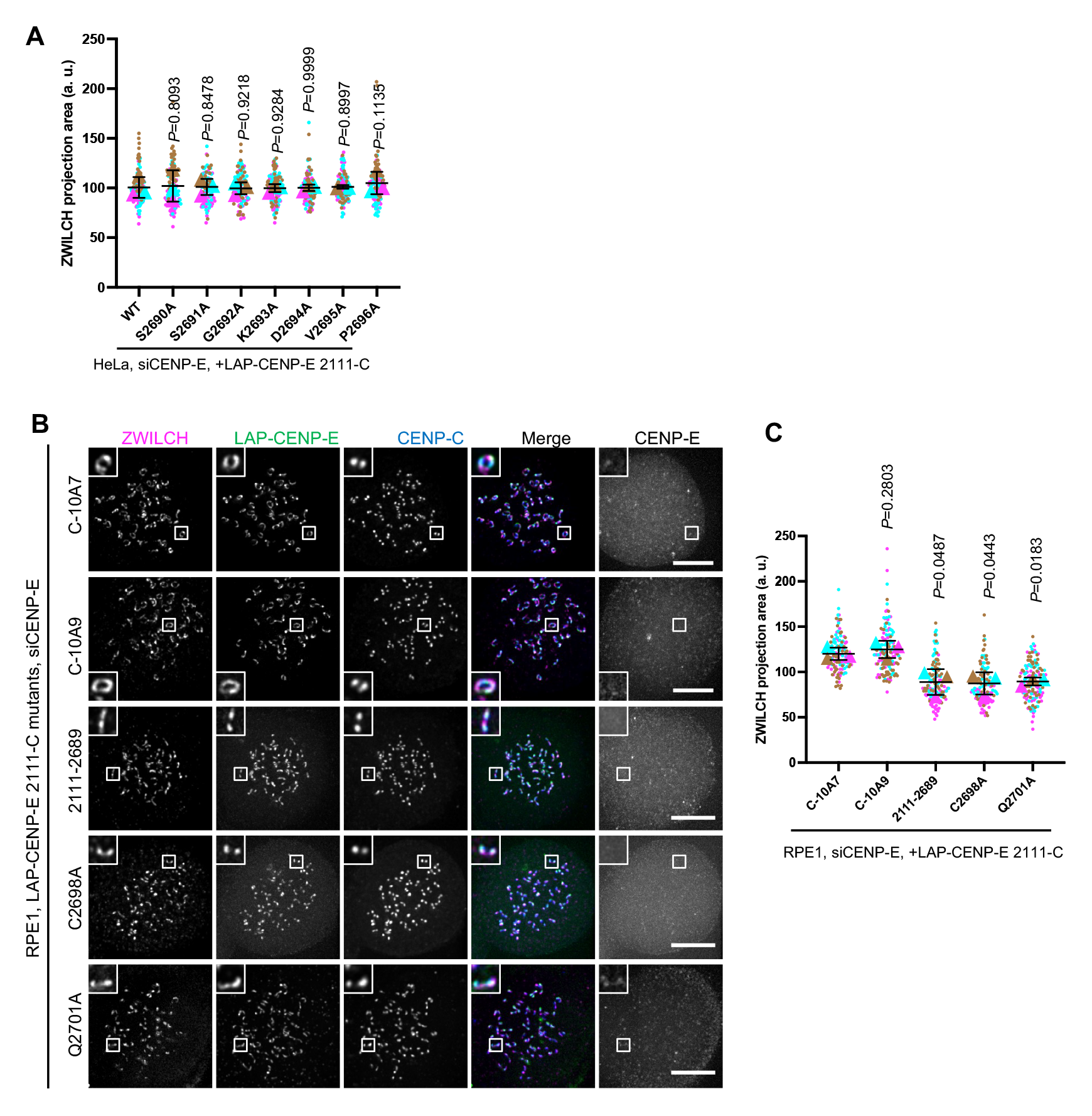
Fibrous corona formation requires C-terminal farnesylation of CENP-E. A: Projection area of ZWILCH in CENP-E-knockdown HeLa cells that over-express the indicated LAP-tagged CENP-E mutant after 8hrs nocodazole treatment. Student’s test. B: Immunostaining of ZWILCH (magenta), CENP-E (gray) and CENPC (blue) in CENP-E-knockdown RPE1 cells that over-express the indicated LAP-tagged CENP-E mutant after nocodazole treatment overnight. Bar, 5 μm. Rabbit polyclonal CENP-E antibody recognizes human CENP-E 955-1571. C: Projection area of ZWILCH in CENP-E-knockdown RPE1 cells that inducibly over-express the indicated LAP-tagged CENP-E mutant with nocodazole treatment overnight. Student’s test.

**Figure S3.**
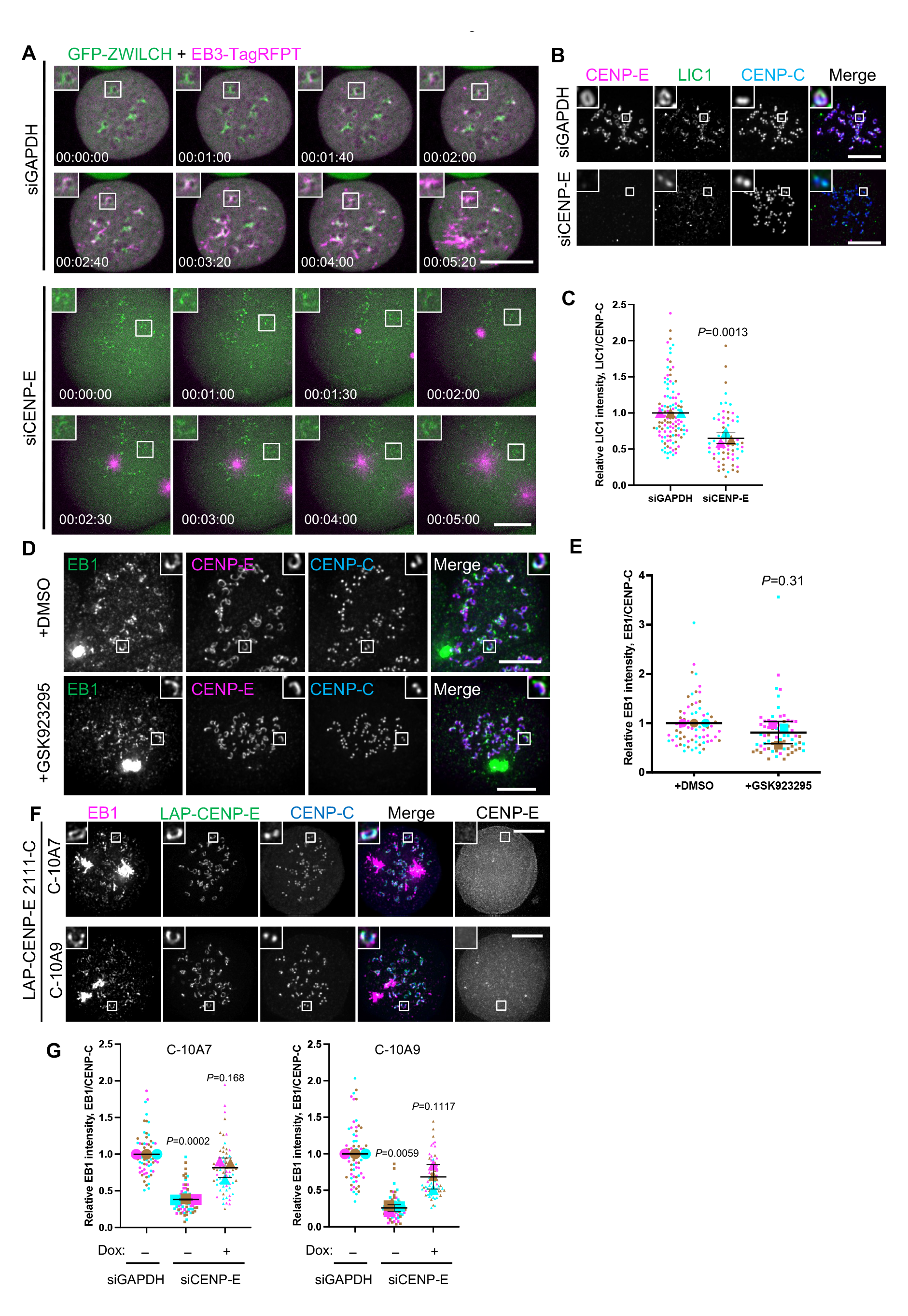
Farnesylated CENP-E is essential for kinetochore-derived microtubule nucleation. A: Stills from live imaging of control and CENP-E-knockdown RPE1 cells that co-express GFP-ZWILCH & EB3-TagRFPT after nocodazole washout. Bar, 5 μm B: Immunostaining of CENP-E (magenta), LIC1 (green) and CENP-C (blue) in control and CENP-E-depleted RPE1 cells treated with nocodazole overnight. Bar, 5 μm. C: Relative LIC1 intensity at the kinetochore compared to CENP-C in control and CENP-E-knockdown RPE1 cells treated with nocodazole overnight. Student’s test. D: Immunostaining of EB1 (green), CENP-E (magenta) and CENP-C (blue) in DMSO and GSK923295-treated RPE1 cells after recovery of 2.5 minutes from nocodazole washout. Bar, 5 μm. E: Relative EB1 intensity at the kinetochore compared to CENP-C in DMSO and GSK923295-treated RPE1 cells after recovery of 2.5 minutes from nocodazole washout. Ratio paired Student’s test. F: Immunostaining of EB1 (magenta), CENP-E (gray) and CENP-C (blue) in CENP-E-knockdown RPE1 cells that over-express the indicated CENP-E mutant after recovery of 3 minutes from nocodazole washout. Bar, 5 μm. Rabbit polyclonal CENP-E antibody recognizes human CENP-E 955-1571. G: Relative EB1 intensity at the kinetochore compared to CENP-C in control and CENP-E-knockdown RPE1 cells that inducibly over-express the indicated CENP-E mutant after recovery of 3minutes from nocodazole washout. Student’s test.

**Figure S4.**
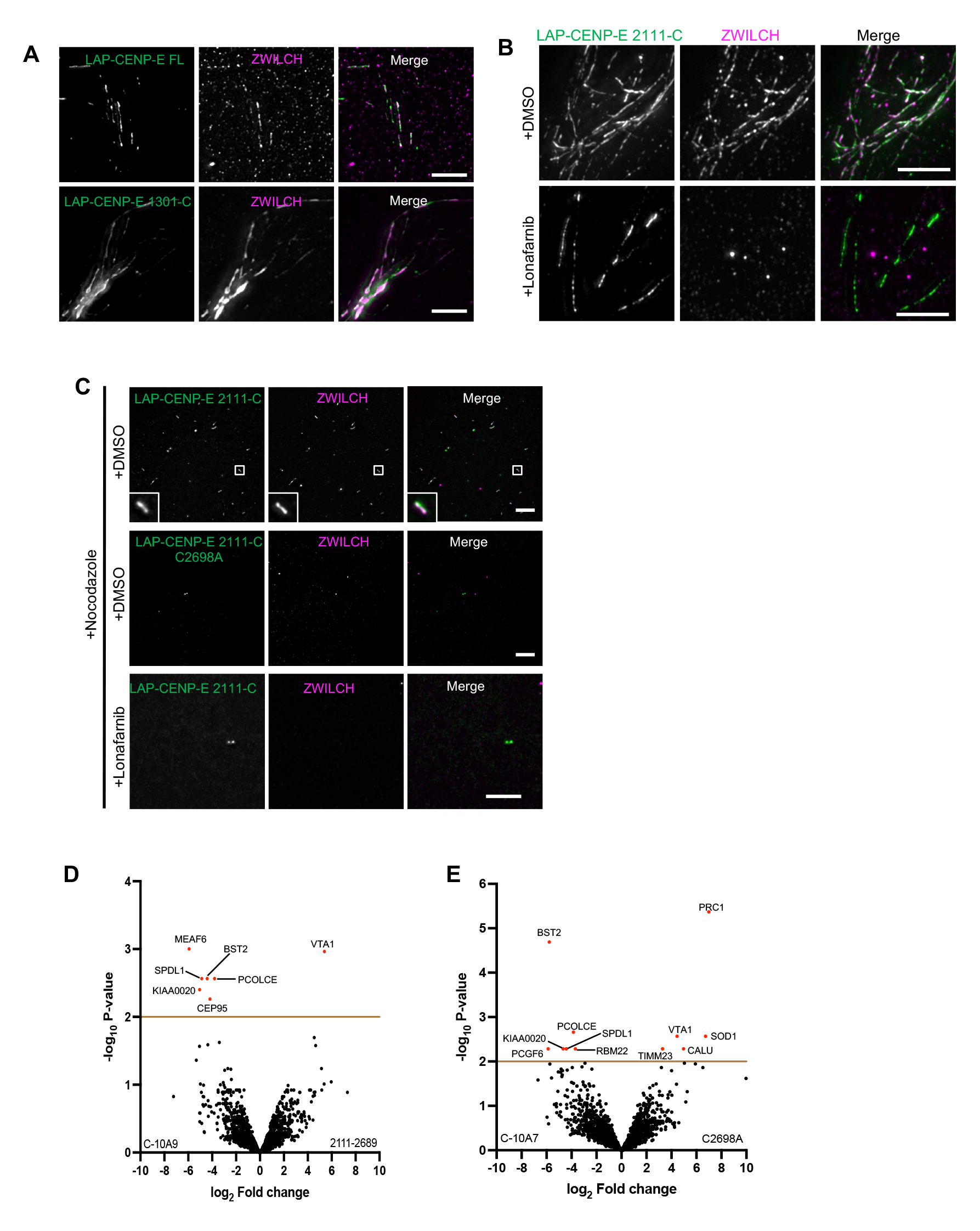
A farnesylated CENP-E fragment can recruit the RZZ-S complex in interphase cells. A: Immunostaining of ZWILCH in RPE cells that over-express LAP-CENP-E FL or LAP-CENP-E 1301-C. B: Immunostaining of ZWILCH (magenta) in control and 5uM lonafarnib-treated HeLa cells that over-express LAP-CENP-E 2111-C. Bar, 5 μm. C: Immunostaining of ZWILCH in DMSO/Lonafarnib-treated RPE cells that over-express LAP-CENP-E 2111-C and LAP-CENP-E 2111-C C2698A in the presence of nocodazole treatment. D: Volcano plot shows top enriched proteins bound to the streptavidin beads incubation with extracts of HeLa cells expressing LAP-TurboID-CENP-E 2111-C C-10A9 (left) or LAP-TurboID-CENP-E 2111-2689 (right). E: Volcano plot shows top enriched proteins bound to the streptavidin beads incubation with extracts of HeLa cells expressing LAP-TurboID-CENP-E 2111-C C-10A7 (left) or LAP-TurboID-CENP-E 2111-C C2698A (right).

**Figure S5.**
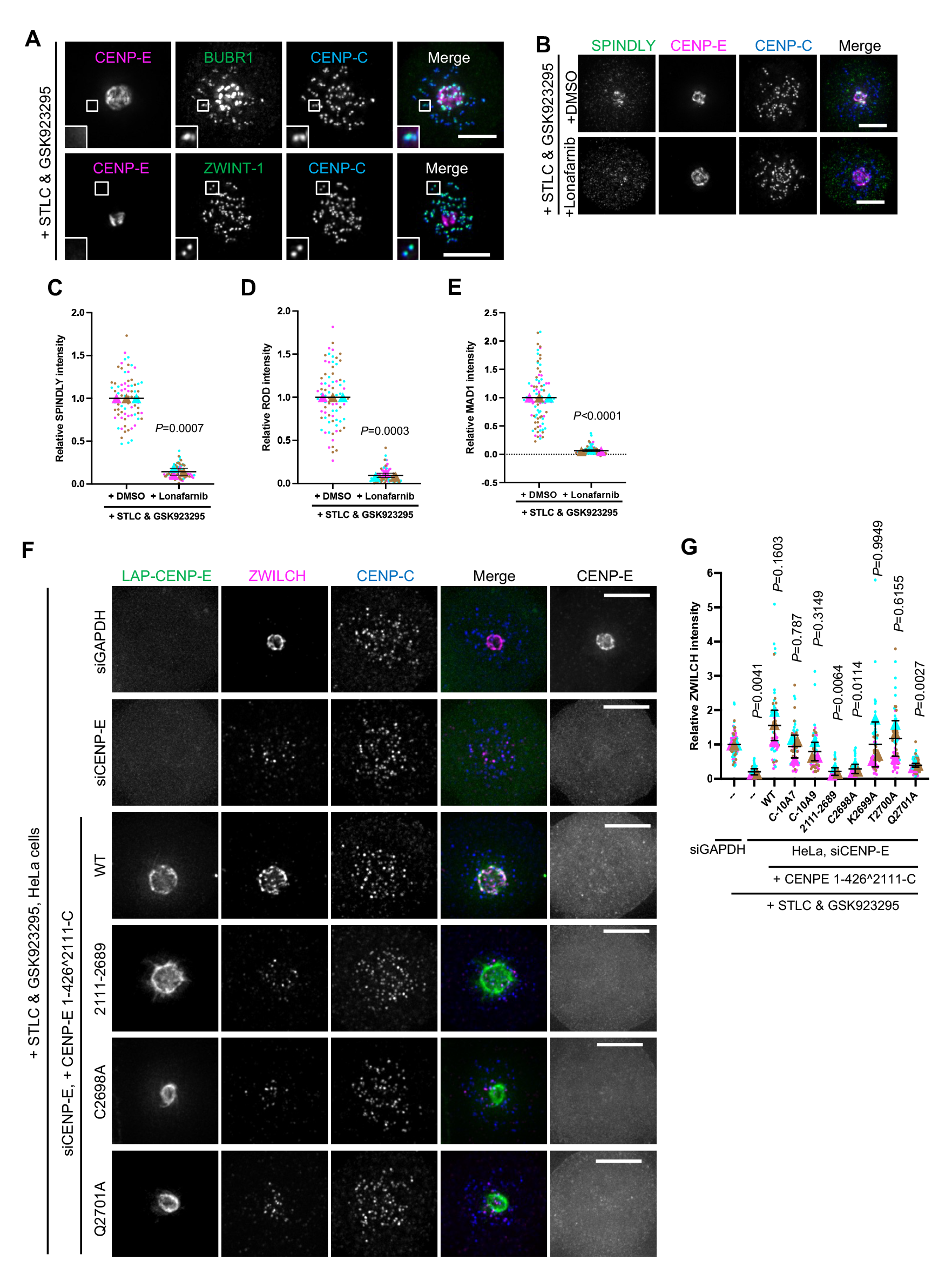
Farnesylation promotes interaction of endogenous CENP-E with fibrous corona proteins in mitosis. A: Immunostaining of BUBR1 (green) / ZWINT-1 (green), CENP-E (magenta) and CENPC (blue) in RPE1 cells treated with STLC and GSK923295 overnight. Bar, 5 μm. B: Immunostaining of SPINDLY (green), CENP-E (magenta) and CENPC (blue) in RPE1 cells treated with STLC and GSK923295 overnight in the presence of DMSO or 5 μM lonafarnib. Bar, 5 μm. C: Relative intensity of SPINDLY at the spindle pole in RPE1 cells treated with STLC and GSK923295 overnight in the presence of DMSO or 5 μM lonafarnib. Student’s test. D: Relative intensity of ROD at the spindle pole in RPE1 cells treated with STLC and GSK923295 overnight in the presence of DMSO or 5 μM lonafarnib. Student’s test. E: Relative intensity of MAD1 at the spindle pole in RPE1 cells treated with STLC and GSK923295 overnight in the presence of DMSO or 5 μM lonafarnib. Student’s test. F: Immunostaining of ZWILCH (magenta), CENP-E (gray) and CENPC (blue) in control and CENP-E-depleted HeLa cells and CENP-E-depleted HeLa cells that over-express the indicated LAP-tagged CENP-E mutant after STLC and GSK923295 treatment overnight. Bar, 5 μm. LAP-tagged CENP-E mutants were generated by fusing the N-terminal CENP-E 1-426 (comprising the motor domain) to the C-terminal CENP-E 2111-C (KT-binding domain and MT-binding domain). Rabbit polyclonal CENP-E antibody recognizes human CENP-E 955- 1571. G: Relative intensity of ZWILCH at the spindle pole in control and CENP-E-depleted HeLa cells and CENP-E-depleted HeLa cells that inducibly over-express the indicated LAP-tagged CENP-E mutant after STLC and GSK923295 treatment overnight. Student’s test.

**Figure S6.**
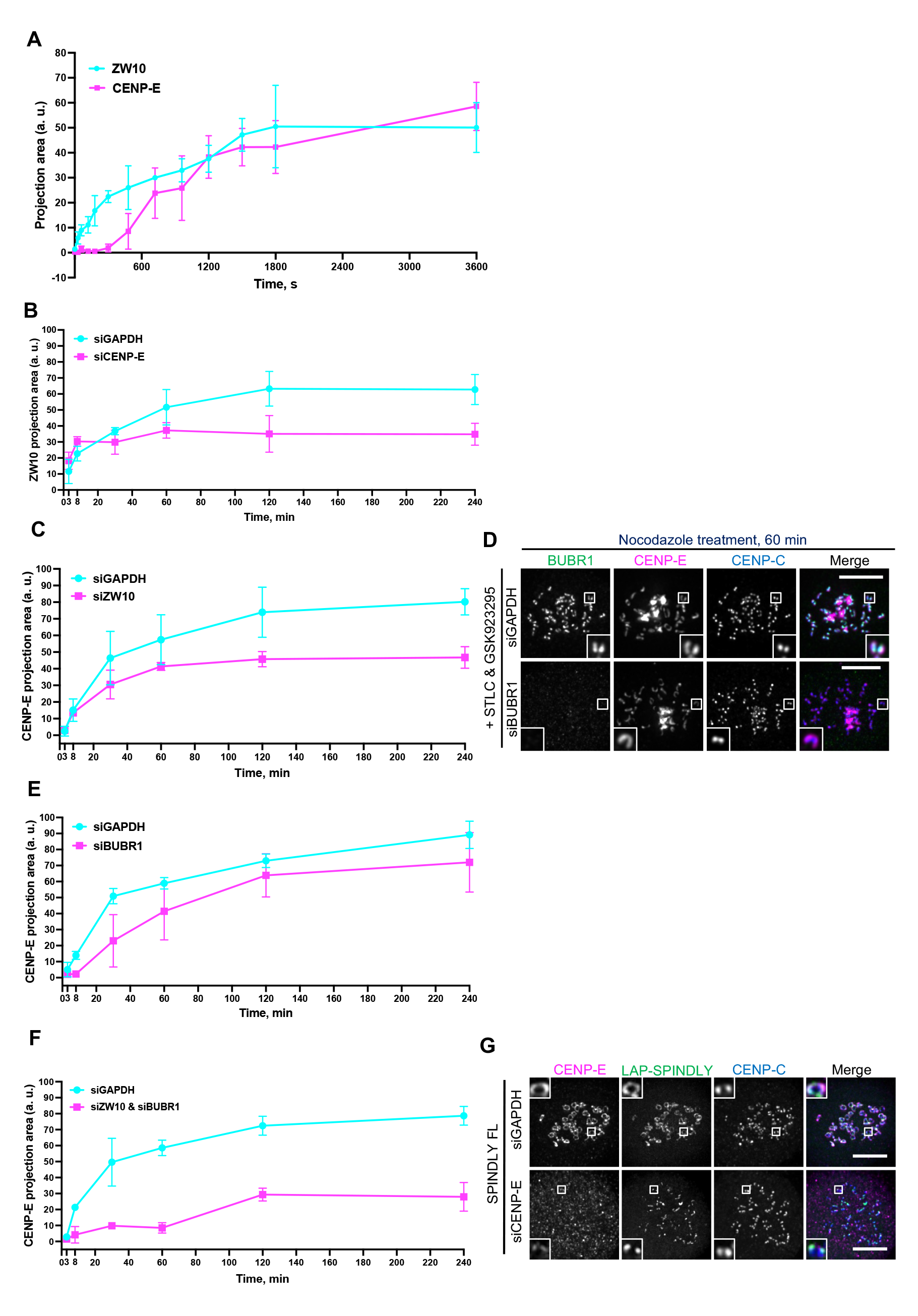
CENP-E impacts fibrous corona formation after initial RZZ-S kinetochore recruitment. A: Projection area of ZW10 and CENP-E in RPE1 cells treated with nocodazole for indicated time, which had been arrested with STLC and GSK923295 overnight. B: Projection area of ZW10 in control and CENP-E-depleted RPE1 cells treated with nocodazole for indicated time, which had been arrested with STLC and GSK923295 overnight. C: Projection area of CENP-E in control and ZW10-depleted RPE1 cells treated with nocodazole for indicated time, which had been arrested with STLC and GSK923295 overnight. D: Immunostaining of BUBR1 (green), CENP-E (magenta) and CENP-C (blue) in control and BUBR1-depleted RPE1 cells treated with nocodazole for indicated time, which had been arrested with STLC and GSK923295 overnight. Bar, 5 μm. E: Projection area of CENP-E in control and BUBR1-depleted RPE1 cells treated with nocodazole for indicated time, which had been arrested with STLC and GSK923295 overnight. F: Projection area of CENP-E in control and ZW10 & BUBR1-co-depleted RPE1 cells treated with nocodazole for indicated time, which had been arrested with STLC and GSK923295 overnight. G: Immunostaining of CENP-E (magenta) and CENP-C (blue) in control and CENP-E-knockdown RPE1 cells that inducibly over-express LAP-SPINDLY FL after nocodazole treatment overnight. Bar, 5 μm.

**Figure S7.**
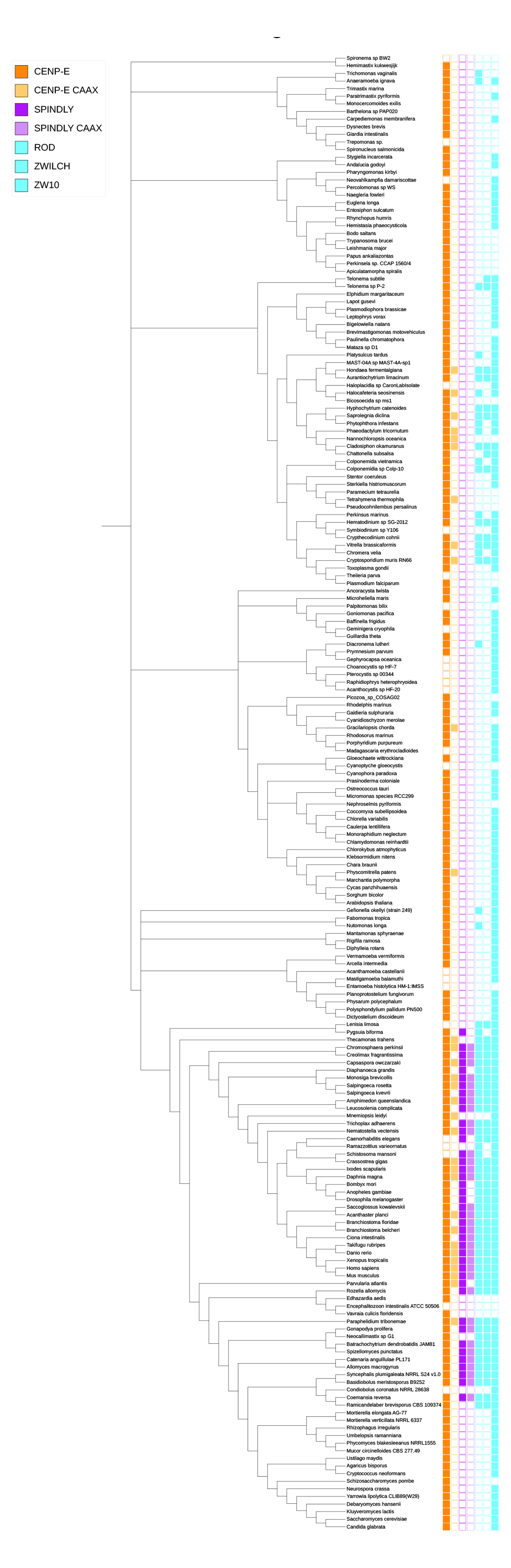
Comparative genomics of fibrous corona formation mechanisms in eukaryotes. Phylogenetic profiling of CENP-E, the CAAX box of CENP-E, SPINDLY, the CAAX box of SPINDLY, ROD, ZWILCH and ZW10 across the entire database representing eukaryotic diversity (194 species). Presences are represented with colored squares, with absences indicated by white squares.

